# SpatialCompassV (SCOMV): *De novo* cell and gene spatial pattern classification and spatially differential gene identification

**DOI:** 10.64898/2026.02.26.707418

**Authors:** Ryosuke Nomura, Shunsuke A. Sakai, Shun-Ichiro Kageyama, Katsuya Tsuchihara, Riu Yamashita

**Affiliations:** Division of Translational Informatics, Exploratory Oncology Research & Clinical Trial Center, National Cancer Center, Kashiwa, Chiba, 277-8577, Japan; Department of Computational Biology and Medical Sciences, Graduate School of Frontier Sciences, The University of Tokyo, Kashiwa, Chiba, 277-8562, Japan; Department of Integrated Biosciences, Graduate School of Frontier Sciences, The University of Tokyo, Kashiwa, Chiba, 277-8563, Japan

## Abstract

Spatial omics technologies enable the detection of gene expression together with spatial information in tissues. However, many existing analytical methods rely on prior biological knowledge or predefined annotations, while being limited in their ability to systematically characterize spatial distribution patterns. Here, we developed SpatialCompassV (SCOMV), a computational tool that clusters genes and cell types based on vectorial relationships between transcript locations and regions of interest, such as tumors. This tool quantifies the spatial positioning of genes and cells relative to a defined reference region by encoding their distance and direction into structured feature representations. SCOMV captured tumor-associated spatial patterns and enabled the unsupervised classification of genes into internal, peripheral, partially peripheral, and ubiquitous distribution types in breast and lung cancer spatial transcriptomic datasets of Xenium. Notably, SCOMV detected immune cell-related signatures that were preferentially localized in CAF-low regions. Extending the analysis to multiple regions of interest further enabled malignant state discrimination. Moreover, SCOMV identifies genes that differ not only in gene expression levels, but also in spatial distribution patterns, which we termed spatially differential genes (spatially DEGs).

## Introduction

The tumor microenvironment (TME), a complex ecosystem composed of tumor, stromal, and immune cells as well as extracellular matrix components, plays a central role in cancer progression, therapeutic response, and immune evasion [1–3]. Single-cell RNA sequencing has become a powerful method for profiling gene expression at single-cell resolution; however, it requires tissue dissociation, which results in the loss of spatial information and prevents *in situ* assessment of immune infiltration, stromal barrier organization, or immune evasion–related processes [4–6]. Recent advances in spatial transcriptomics (ST) technologies, such as CosMx [7], Visium [8] and Xenium [9], have overcome this limitation by measuring gene expression across tissue sections while maintaining spatial coordinates, thereby enabling *in situ* investigation of cellular organization and cell–cell interactions within the TME [10–12].

Although ST is a powerful technology, extracting biologically meaningful information from ST data remains challenging because of its high dimensionality [13, 14]. Previous approaches have often relied on manual inspection of spatial expression for hundreds or thousands of genes, making the process extremely time-consuming and susceptible to observer bias. To overcome this limitation, computational frameworks, such as Scanpy [15] and STAGATE [16] have been developed to automate cell-based clustering, annotation, and graph-based embedding for spatial transcriptomic analysis. In parallel, methods such as Seurat [17] and Squidpy [18] identify spatially varying genes (SVGs) by quantifying spatial dependencies and nonrandom spatial structures using spatial autocorrelation metrics such as Moran’s I and Geary’s C [19, 20]. These metrics are well suited for detecting broad spatial trends in gene expression and capturing large-scale structures across tissue sections.

Despite these advances, the current approaches still face two major limitations when applied to tumor tissue analysis. First, existing methods are not well-suited for classifying gene distributions relative to standard regions, such as tumor areas or other structural landmarks. Consequently, distinguishing whether transcripts are expressed within the tumor, at the tumor-stromal boundary, or in peripheral stromal compartments is difficult. Even immune-related genes that are frequently examined in spatial analyses exhibit highly diverse distribution patterns (Supplementary Fig. S1) [21–24]. Immune checkpoint–associated genes, for example, show heterogeneous spatial localization: TIGIT and CD274 display distinct enrichment patterns at tumor boundaries, highlighting the difficulty of systematically categorizing such features using current approaches [25, 26]. Second, many current methods rely on static and predefined databases. Tools such as Squidpy [18] and NATMI [27] quantify cell–cell interactions using curated ligand–receptor pairs, whereas others depend on marker-driven cell-type annotations derived from single-cell or bulk transcriptomic data. As the scale of ST increases, data-driven *de novo* frameworks that can construct new knowledge bases for integrating spatial interaction patterns are increasingly required.

In this study, we developed ‘SpatialCompassV’ (SCOMV), a computational tool that characterizes gene localization relative to a defined reference region (the tumor in this case) into several distinct spatial patterns, including internal (localized within tumor regions), peripheral (enriched at tumor boundaries), partially peripheral (localized to only a subset of boundary regions), ubiquitous (distributed broadly across the tissue), and combinations of these patterns. The tool supports downstream analyses, including (1) classification of gene distribution patterns, (2) inference of cell–cell and gene–gene interactions, and (3) estimation of tumor-associated malignancy signatures from spatial patterns in a manner that does not rely on predefined biological annotations. Collectively, these capabilities enable the systematic interpretation of tumor-related gene localization patterns, particularly peripheral and partially peripheral expression patterns relevant to tumor–immune interactions.

## Materials and methods

### Signed minimum distance vector (SMDV) derivation

For each grid location, we computed the shortest Euclidean vector for the nearest grid belonging to the tumor region. When multiple nearest tumor grids were present, all candidate vectors were retained. These vectors were then reoriented by inverting their direction such that they consistently pointed outward from the tumor boundary. To distinguish whether a grid was located inside or outside the tumor region, the vector magnitude was treated as a signed distance; grids inside the tumor region were assigned negative values, whereas those outside the tumor region retained positive values. Each resulting vector is defined as a signed minimum distance vector (SMDV).

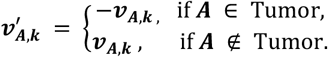

Next, for each grid, the expression level of each gene was obtained and denoted as *E*_*g,i*_, where *g* represents the gene, and *i* represents the grid index. For a given gene *g*, the vector from grid *A* to the tumor region, denoted as *V*_*g,A*_, is defined as follows:

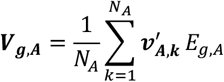

where 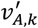 is the signed shortest vector defined above, and *N*_*A*_ is the number of candidate shortest paths for grid *A*. To obtain the overall spatial representation for gene *g*, the above equation was performed for all grids *A* ∈ 𝒢, yielding the complete set of gene-specific vectors:

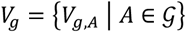

where 𝒢 denotes the set of all grids within the analyzed field of view.

### Polar coordinate map construction

For each gene (or cell), a polar coordinate map was generated by dividing the space into bins of 10 μm in radius and 30° in angle. Each bin is assigned the corresponding vector *V*_*g,A*_ based on its spatial position. All binned vectors were normalized such that their total sum was equal to one.

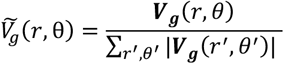

### Calculation of distribution similarity and similarity matrix

For any two genes *g*_*i*_ and *g*_*j*_, their spatial distributions were represented as normalized polar coordinate maps, 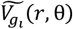 and 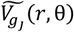 constructed using the same predefined (*r*, θ) binning scheme. For each corresponding (*r*, θ) bin, the minimum of the two values was computed and summed across all bins to quantify spatial similarity:

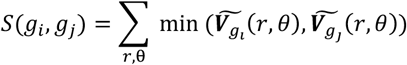

This value reflects the degree of overlap between two spatial distributions, ranging from 0 (no overlap) to 1 (identical distributions). By computing *S* (*g*_*i*_, *g*_*j*_) for all gene pairs, similarity matrix *S* was constructed as follows:

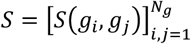

where *N*_*g*_ denotes the total number of genes analyzed. Calculating *S* (*g*_*i*_, *g*_*j*_) for all the gene pairs yielded the similarity matrix *S* = [*S* (*g*_*i*_, *g*_*j*_)]

### Clustering and heatmap visualization of cellular distribution

For the similarity matrix representing the spatial distribution of cells, pairwise dissimilarity was defined as *D*(*i, j*) = 1 − *S*(*i, j*), where *S*(*i, j*) denotes the similarity between cells *i* and *j*. Hierarchical clustering was performed to compare cellular spatial patterns. Hierarchical clustering was performed using Euclidean distance with Ward’s linkage, implemented in the Python library seaborn (v0.13.2; ‘clustermap’ function). The resulting dendrogram and heatmap visualized groups of cells that exhibited similar spatial distribution patterns.

### Principal coordinate analysis (PCoA) of gene distribution

PCoA was applied to the gene-based similarity matrix to visualize the global relationships among genes according to their spatial distributions. The distance matrix was subjected to eigen decomposition to obtain the principal coordinate axes. The analysis was performed using the Python library scikit-bio (v0.7.0; ‘pcoa’ function), and the first and second principal coordinates were used for visualization. The resulting PCoA coordinates were subsequently used to cluster and interpret the groups of genes with similar spatial distribution patterns.

### Symmetric non-negative matrix factorization (SymNMF)

For each region of interest (ROI), the similarity matrix 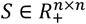 was decomposed using symmetric non-negative matrix factorization (SymNMF) [28]. SymNMF approximates ***S*** as the product of a low-rank nonnegative matrix ***W*** and its transpose:

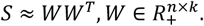

The objective function was defined as:

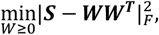

where | ⋅ |_F_ denotes the Frobenius norm and *k* represents the number of components (here, *k* = 1). Factor matrix ***W*** was optimized using standard multiplicative update rules. The iterations were repeated for a maximum of 500 times. These processes were implemented using the publicly available code from the ‘dakuang/symnmf’ GitHub repository (https://github.com/dakuang/symnmf).

The resulting component vectors were averaged across the columns to obtain 1-dimensional feature vectors. These vectors were hierarchically clustered using Ward’s linkage with Euclidean distance, implemented in the Python library SciPy (v1.11.4; ‘scipy.cluster.hierarchy’ module). Subsequently, principal component analysis (PCA) was performed on these vectors, and the first 20 principal components were used to evaluate the contribution of each gene to the clustering structure. Genes with the highest and lowest contributions were extracted as spatially differentially expressed genes (spatial DEGs), which were used to characterize spatial expression patterns across multiple tumor regions.

### Data acquisition

Breast cancer data from the Xenium platform were obtained from a public repository (https://www.10xgenomics.com/jp/products/xenium-in-situ/preview-dataset-human-breast). The ReadXenium function from the Python library stlearn (v0.4.12) was used to read the hematoxylin and eosin (H&E) images (https://www.dropbox.com/s/th6tqqgbv27o3fk/CS1384\_post-CS0\_H\%26E\_S1A\_RGB-shlee-crop.png?dl=1) and the accompanying gene expression and cell coordinate files (Xenium_FFPE_Human_Breast_Cancer_Rep1_cell_feature_matrix.h5 and Xenium_FFPE_Human_Breast_Cancer_Rep1_cells.csv.gz). The lung cancer data from the Xenium platform were obtained from previous patients [29]. This study was approved by the Institutional Review Board of the National Comprehensive Cancer Study Board (NCCHE) Institutional Review Board (Protocol Number 2022-407).

### Gene filtering

Genes exhibiting uniformly high expression across most cells (e.g., housekeeping-like genes, which are constitutively expressed to maintain basic cellular functions; expression > 95th percentile within each field of view) were excluded, as they are generally not informative for spatial pattern analysis and increase computational burden without contributing meaningful spatial information [30, 31]. Genes with notably low overall expression (< 35th percentile) were excluded to avoid noisy distributions arising from sparse detection. In contrast, genes with low global expression but positive spatial autocorrelation (Morab’s I ≥ 0) were retained, as locally concentrated enrichment may reflect biologically meaningful events.

### Definition of regions of interest (ROIs)

In the breast cancer panel, ten ROIs were defined as rectangular areas on the image plane using image column (x) and image row (y) coordinates. Each ROI was specified as follows:

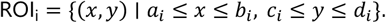

ROIs 1–6 correspond to ductal carcinoma in situ (DCIS) and ROIs 7–10 correspond to invasive ductal carcinoma (IDC). We selected ROI6 for analysis of Fig. 2 and Fig. 3. The coordinate ranges (*a*_*i*_, *b*_*i*_, *c*_*i*_, *d*_*i*_) for each ROI are presented in Supplementary Table S1. For the lung cancer panel, analyses were performed using a single ROI defined using the same coordinate-based formulation described above.

## Results

### Overview of SCOMV

We developed SCOMV, a computational tool designed to quantify spatial distribution patterns at both the gene and cell levels relative to tumor regions, for characterizing the spatial relationships between tumors and surrounding cells. Using SCOMV, the spatial distribution patterns were categorized into four representative types (Fig. 1A):

i. Internal, primarily expressed within the tumor.
ii. Peripheral, circumferentially distributed around the tumor.
iii. Partially peripheral, restricted to specific segments of the tumor boundary.
iv. Ubiquitous, broadly distributed across the tissue.

SCOMV consists of three main steps as below:

1. Tumor segmentation and vectorization: Spatial data were partitioned into grid units, and tumor regions were identified using Spatial Knife Y (SKNY) [32] (Fig. 1B). The signed minimum distance vectors (SMDVs) from each grid to the nearest tumor cell were calculated and scaled according to the analytical focus, using cell counts for cell-level analyses and gene expression levels for gene-level analyses, thereby emphasizing the contributions from abundant cells or genes in the overall distribution pattern.
2. Spatial feature embedding and distribution clustering for cells and genes: SMDVs were transformed into polar coordinates, which were then used to compute pairwise similarities between cells and genes (Fig. 1C). The resulting similarity matrices enabled downstream analyses to group spatial distribution patterns, with cell-level similarities summarized using heatmap-based visualization and gene-level pattern clustering performed using PCoA.
3. Unsupervised clustering of tumor regions via latent gene distribution pattern decomposition: Gene–gene similarity matrices derived from multiple tumor regions were decomposed using symmetric non-negative matrix factorization (SymNMF) to extract latent gene distribution patterns shared across regions (Fig. 1D). The tumor regions were classified in an unsupervised manner based on these latent features. Gene contribution scores derived from these clusters identified key genes associated with distinct tumor microenvironmental characteristics.

**Figure 1.**
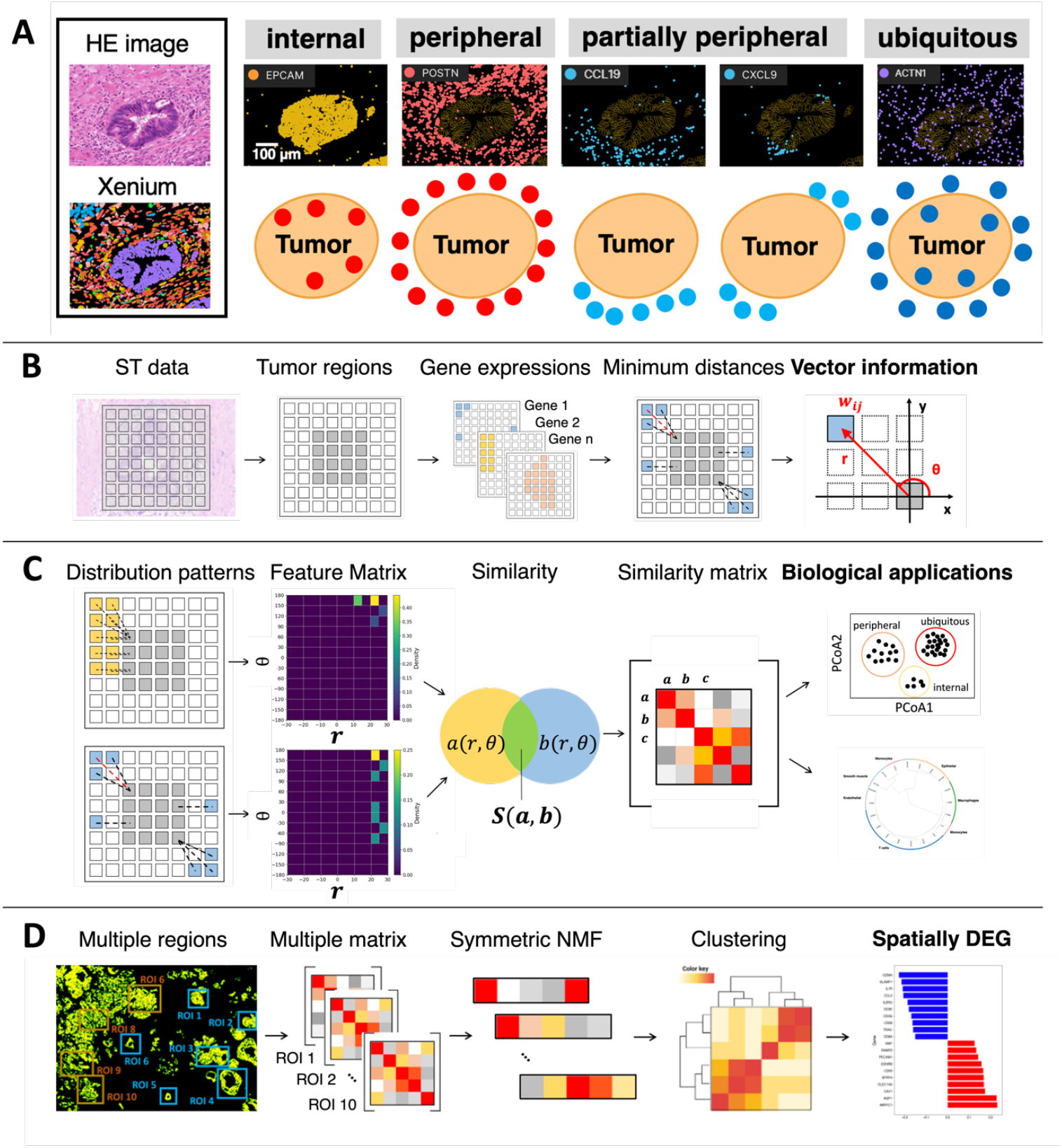
SCOMV algorithm overview. (A) Gene distribution patterns relative to the tumor region. (B) Workflow for calculating vector-based spatial features. Gene expression data and tumor regions were imported via AnnData and processed using SpatialKnifeY. The SMDV from each grid to the nearest tumor grid was calculated, weighted by expression levels. (C) Comparative analysis of spatial distribution patterns. SMDVs were transformed into polar coordinates, converted to feature maps, and compared across genes to construct a similarity matrix for clustering. (D) Multiple region integration. Dimensionality reduction and clustering of similarity matrices from multiple fields identified spatially DEG.

### Clustering of cell distribution patterns

We first delineated the tumor region using SKNY to define the tumor boundaries and subsequently selected a region of interest (ROI). For both the SKNY-derived tumor boundaries and the selected ROI, the spatial extent showed close agreement with the corresponding H&E-stained sections (Fig. 2A–B), indicating that the regions used in subsequent analyses were consistent with the underlying tumor morphology (Supplementary Table S2).

**Figure 2.**
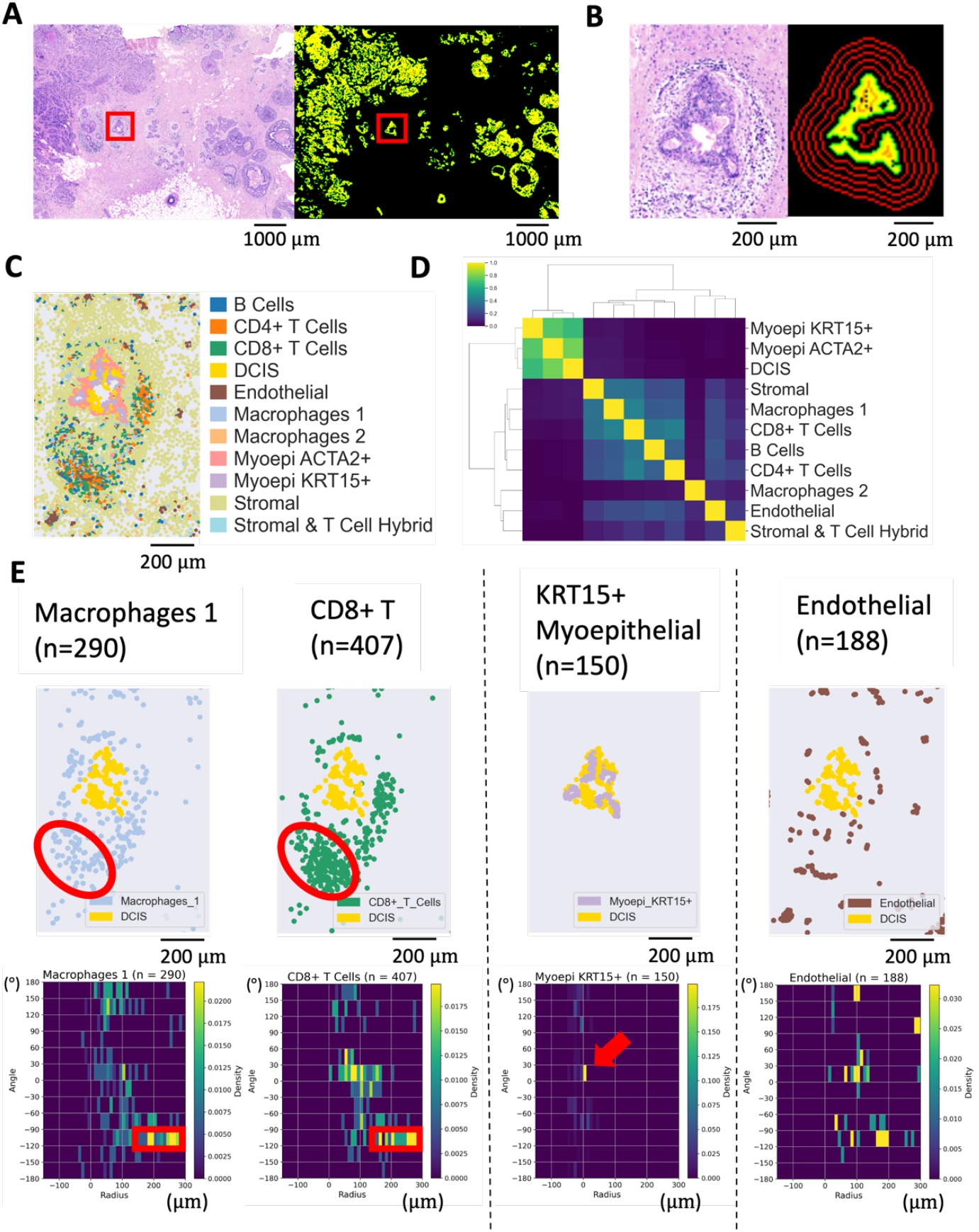
Cell distribution analysis in breast cancer sample. (A) Hematoxylin and eosin (H&E) stain (left) and tumor regions extracted by SpatialKnifeY (right), with the analyzed region of interest (ROI, red box). (B) Enlarged view of the selected ROI. (C) Spatial distribution of annotated cell types across the ROI. (D) Heatmap and hierarchical clustering based on similarities in cell distribution patterns. (E) Representative spatial distributions and corresponding polar coordinate maps for each cell type. In the polar plots, the radial axis denotes distance (−150–300 µm), and the angular axis denotes angle (−180°–180°).

To assess whether SCOMV could recapitulate previously reported spatial localization patterns of annotated cell types, we used previously published annotations to generate signed minimum distance vector (SMDV)-based polar coordinate maps for each cell type [9]. The pairwise similarities among these maps were used to construct a similarity matrix, which was subsequently subjected to hierarchical clustering. Cluster analysis identified distinct clusters of tumor-associated cell types (Fig. 2D and Supplementary Table S3). M1 macrophages, CD4^+^ and CD8^+^ T cells, and B cells were grouped into a single cluster that was preferentially localized at distances of approximately 200–300 µm from the tumor boundary and at angular positions between −120° and −90° (Fig. 2E, left and Supplementary Fig. 2, 3). In contrast, myoepithelial cells formed a separate cluster and were l predominantly localized near 0 µm, corresponding to intra-tumoral regions (Fig. 2E, center and Supplementary Fig. 4). Endothelial cell and stromal/T-cell hybrid populations displayed more dispersed spatial patterns, characterized by multiple weaker peaks across the tissue (Fig. 2E, right and Supplementary Fig. 5). Collectively, these observations suggest that SCOMV captures the characteristic positional features of different cell types relative to tumor regions.

### Clustering of gene distribution patterns

Having demonstrated that SCOMV captures cell-level spatial distribution patterns relative to tumor regions, we examined the spatial distribution patterns of individual genes to characterize gene-specific localization relative to tumor regions. SMDVs were calculated and pairwise similarities among representative genes were quantified. Visual inspection of the Xenium dataset suggested that the genes exhibited four previously defined tumor-related localization patterns: internal, peripheral, partially peripheral, and ubiquitous. The resulting gene–gene similarity matrix was projected into a two-dimensional space using PCoA, and representative cell-type marker genes were annotated on the PCoA plot for reference (Fig. 3A and Supplementary Table S4). The PCoA plot showed several distinct gene clusters, with genes sharing similar spatial localizations and tending to occupy nearby positions in the PCoA space (Fig. 3B–C). For example, epithelial cell markers such as KRT8 and EPCAM, which exhibited predominantly internal localization, were positioned toward the low end of the PCoA1 axis (e.g., KRT8: PCoA1 = −0.518; EPCAM: PCoA1 = −0.435). In contrast, smooth muscle-associated genes, including ACTA2 and ACTG2, which exhibited both intra-tumoral and peripheral distribution pattern, occupied intermediate positions along the PCoA1 axis (e.g., ACTA2: PCoA1 = −0.151; ACTG2: PCoA1 = −0.0548). Immune-related genes, such as CD68 and CD8A, were distributed toward high values along the PCoA1 axis (e.g., CD68: PCoA1 = 0.218; CD8A: PCoA1 = 0.255), consistent with their peripheral localization relative to the tumor boundary. Within the immune-related group, macrophage- and T-cell–associated genes were separated into distinct subregions along the PCoA1 axis, suggesting a cell-type-specific spatial organization.

**Figure 3.**
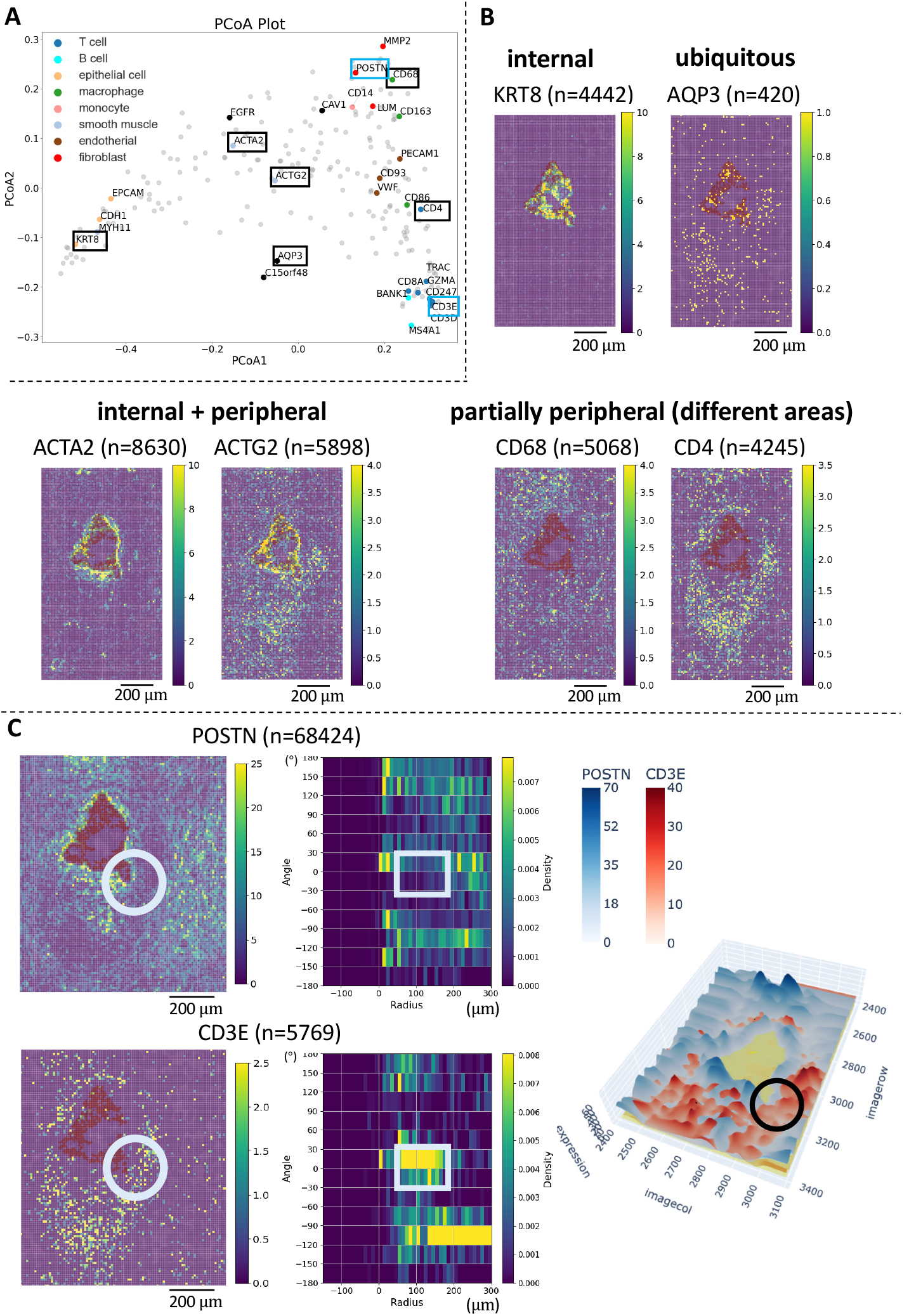
Gene distribution analysis relative to tumor regions in breast cancer panel. (A) Principal coordinates analysis (PCoA) based on the similarity matrix of gene distribution patterns, where each point represents a gene. (B) Spatial expression patterns of representative genes selected from PCoA space. (C) Distributions and polar coordinate maps of POSTN and CD3E (left), illustrating their spatial relationship (right; tumor region, yellow; POSTN, blue; CD3E, red). Color bars indicate expression values scaled to a maximum of 100 for each gene and smoothed using a Gaussian kernel.

We compared our method with the existing spatial transcriptomic analysis approaches. Moran’s I and Geary’s C scores were computed for all genes, and the spatial expression patterns of the top-ranked genes were examined (Supplementary Fig. S7, 8 and Supplementary Table S5, 6). These methods identify genes with highly clustered expression; however, the top-ranked genes represent a mixture of internal, peripheral, and ubiquitous localization patterns, limiting their ability to provide an ordered representation based on the tumor-relative spatial distribution.

POSTN is expressed in cancer-associated fibroblasts (CAFs) surrounding tumor regions and contributes to tumor progression, although its precise spatial distribution relative to the tumor regions remains incompletely characterized [33, 34]. In the PCoA projection, POSTN occupied a position distinct from that of the immune-related gene clusters (Fig. 3A, blue square). Its spatial distribution showed a predominantly peripheral pattern, with notably reduced expression in the lower-right portion of the tumor region (Fig. 3C, upper-left, circle). In contrast, the immune cell marker CD3E showed high expression around this region, a pattern observed in both the raw spatial distribution (Fig. 3C, lower left, circle) and the corresponding polar coordinate map (Fig. 3C, lower left, square). Taken together, these observations suggest that SCOMV captures localized features of the tumor microenvironment that are not readily discernible from conventional spatial distributions.

### Applying SCOMV to a lung cancer dataset

Next, we evaluated SCOMV using a non–small cell lung cancer (NSCLC) dataset to assess its applicability to different cancer types. The PCoA coordinates derived from the gene–gene similarity matrix showed spatial gene distribution patterns that were highly consistent with those observed in breast cancer (Supplementary Fig. S9 and Supplementary Table S7, 8, 9). Specifically, the epithelial, stromal, and immune-related gene groups formed distinct clusters in the PCoA space. Genes such as MKI67, which are predominantly expressed near the cell membrane, formed a separate cluster from these groups. These results suggest that SCOMV is applicable to different tumor types.

### Integrated analysis across multiple fields and spatial DEG identification

In the preceding analyses, we classified gene distribution patterns using a gene–gene similarity matrix constructed from pairwise similarities of gene distribution patterns relative to tumor regions. We hypothesized that this similarity matrix captures compact representations of the tumor microenvironment, such that comparisons across samples may facilitate the unsupervised classification of tumors based on spatial patterns associated with malignancy. To evaluate this possibility, we applied SCOMV to ten regions of interest (ROIs) from a breast cancer dataset, including six ductal carcinoma in situ (DCIS) and four invasive ductal carcinoma (IDC) regions (Fig. 4A). The analysis was restricted to 157 genes that were consistently detected across all the ROIs. For each ROI, SCOMV generated a tumor-centric similarity matrix, which was subsequently subjected to dimensionality reduction using non-negative matrix factorization (NMF), followed by hierarchical clustering. The ten ROIs were separated into three major clusters, and the resulting dendrogram distinguished DCIS from IDC (Fig. 4B).

**Figure 4.**
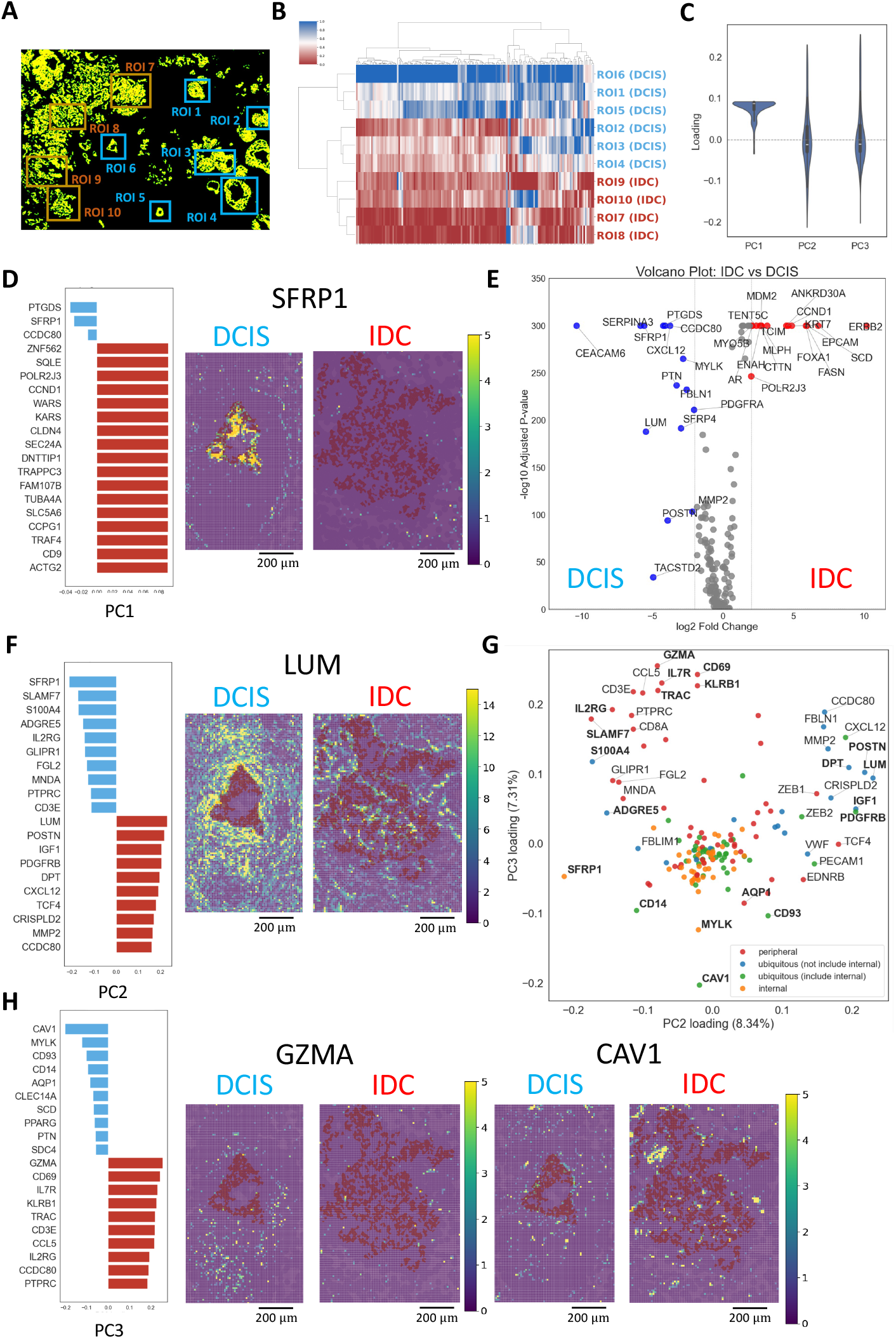
Multiple regions analysis and spatial DEG identification. (A) Ten regions of interest (ROIs) selected from the breast cancer dataset. (B) Hierarchical clustering of similarity matrices constructed from gene distribution patterns in each ROI. (C) Violin plots of PC1, PC2, and PC3. (D) Uppers (red) and lower (blue) values of PC1 and spatial distribution of SFRP1 in DCIS and IDC. (E) Differential gene expression analysis between IDC and DCIS. (F) Top and bottom 10 loadings of PC2 and the spatial distribution of LUM. (G) PC2–PC3 two-dimensional plot. Colors denote predefined spatial distribution patterns: red, peripheral; blue, ubiquitous (excluding internal); green, ubiquitous (including internal); orange, internal. Gene sets ranked in the top five loadings of PC1–PC3 are shown in bold, and those within the top ten in regular font. Internal and ubiquitous (including internal) gene sets cluster in the center. (H) Top and bottom 10 loadings of PC3 and spatial distributions of GZMA and CAV1.

PCA was performed on 157 clustered genes to identify genes contributing to the separation of DCIS and IDC. PC1 showed a narrow distribution, with similar values across most genes, whereas PC2 and PC3 exhibited greater variability (Fig. 4C and Supplementary Table S10). PC1 explained 73.4% of the total variance, whereas PC2 and PC3 explained 8.34% and 7.31%, respectively. Only three genes, SFRP1, PTGDS and CCDC80, had negative PC1 values (Fig. 4D, left). Examination of their spatial distribution indicated that these genes were rarely detected in IDC, whereas they remained detectable in the peripheral regions of DCIS (Fig. 4D, right and Supplementary Fig. 23-25). Consistent with this pattern, these genes were also significantly upregulated in DCIS by differential expression gene (DEG) analysis comparing DCIS and IDC (Fig. 4E). In contrast, other DCIS-enriched genes, such as ZNF562, retained detectable expression in IDC (Supplementary Fig. 26). These results suggest that PC1 primarily reflects the presence or absence of gene expression within tumor-containing regions.

Next, we investigated PC2. Among the top ten positive loads, CAF-associated genes, including LUM and POSTN, were enriched (Fig. 4F, left). These genes are highly expressed in DCIS and predominantly show peripheral spatial localization (Supplementary Fig. 27-30).When PC2 and PC3 were plotted in a two-dimensional space, these CAF-related genes formed a distinct cluster (Fig. 4G, right) and were enriched on the DCIS side of the DEG analysis (Fig. 4E and Supplementary Table S11).

PC3 highlights a distinct set of genes with opposing loading directions. Genes with high positive loadings included immune-related genes, such as GZMA and CD3E (Fig. 4H, left), which were highly expressed in DCIS and displayed partial peripheral localization patterns. Conversely, genes with strong negative loadings, including CAV1 and AQP1, which are associated with vascular or endothelial compartments, were enriched in DCIS but exhibited mixed internal and peripheral distributions (Fig. 31-36). These immune- and vascular-related gene sets form separate clusters in the PC2–PC3 two-dimensional space (Fig. 4G, upper left and lower center, respectively).

Collectively, these analyses indicate that SCOMV captures the differences in spatial gene organization between DCIS and IDC. The genes contributing to this separation differed both in expression levels and spatial distribution profiles, suggesting that a subset of genes can be regarded as “spatially differentially expressed genes”.

## Discussion

In this study, we developed SCOMV, a computational tool that characterizes tumor-relative spatial organization by encoding the distance and angular information. We demonstrated that SCOMV enables the classification of cell-type distribution patterns relative to tumor regions at the cell level. (Fig. 2A-E). Extending this tool to the gene level allows for the unsupervised classification of genes into internal, peripheral, and partially peripheral distribution patterns. Internal genes were predominantly associated with epithelial programs, whereas peripheral genes were enriched for immune- and cancer-associated fibroblast– related functions (Fig. 3A). Notably, these spatial patterns cannot be distinguished using conventional spatial metrics such as Moran’s I or Geary’s C [19, 20]. These statistics summarize spatial autocorrelation as a single scalar value per gene, and therefore do not support direct comparisons of spatial distribution patterns across genes. Consequently, genes with similar metric values may exhibit markedly different spatial organization, such as internal versus peripheral localization (Supplementary Fig. S7, 8, 11, 12). In contrast, SCOMV represents each gene using a spatial feature map and clusters genes based on tumor-relative pattern similarity relative to a reference region, enabling the classification of distribution types with respect to standard regions.

The application of SCOMV to breast and lung cancer datasets revealed consistent tumor-relative spatial distribution patterns across disease contexts. In breast cancer, both POSTN (CAF-related genes) and CD3E (immune-related genes) exhibit peripheral localization relative to the tumor regions, with POSTN being preferentially enriched in peripheral areas where CD3E signals are relatively attenuated (Fig. 3C)[35]. Stromal remodeling and CAF-rich microenvironments can restrict immune–tumor interactions by modulating immune cell activity and reducing physical proximity [36–38]. Although stromal alterations in ductal carcinoma in situ (DCIS) are associated with limited immune cell infiltration, lesions exhibiting histopathological features of tumor regression demonstrate increased immune cell evolvement [39–41]. Taken together, these observations support the possibility that immune cells may preferentially engage tumor regions where CAF-mediated physical or functional barriers are diminished, particularly in areas with reduced POSTN expression in the CAF-rich stroma. In lung cancer, chemokines, such as CXCL9 and CXCL13, showed spatial distributions distinct from those of general T cell markers (Supplementary Fig. S9), consistent with previous observations of preferential accumulation in peritumoral regions following immune checkpoint–based therapies [42, 43]. Although methods such as stLearn and Squidpy can infer cell–cell interactions, these approaches rely on predefined ligand–receptor databases and typically summarize interactions as global measures within a region rather than resolving region-specific spatial organization [18, 44]. In contrast, SCOMV enables data-driven detection of localized interaction patterns without prior biological assumptions by modelling spatial relationships in terms of distance and angular orientation, thereby allowing the identification of confined associations, such as those between CD3E and POSTN.

By extending the analysis across multiple regions of interest, SCOMV further enabled unsupervised tumor region clustering according to malignancy (Fig. 4B) and identification of spatially differential genes (Fig. 4D, F, H). Conventional differential expression gene (DEG) analysis, which relies on differences in gene expression levels without incorporating spatial information, identified several genes enriched in IDC, including epithelial markers such as CDH1 and EPCAM (Fig. 4E) [45, 46]. However, these genes predominantly exhibited internal distribution patterns in both DCIS and IDC, indicating that the observed DEG signals primarily reflected differences in tumor size rather than changes in spatial organization (Supplementary Fig. S37-39). Similarly, genes enriched in DCIS exhibited heterogeneous tumor-related distribution patterns, including internal (SFRP1) and peripheral (LUM) localizations (Fig. 4D, E, F). By integrating expression abundance with tumor-relative spatial positioning, SCOMV distinguished genes that differed only in their expression levels from those that showed distinct distribution patterns (Fig. 4D, F, H). Notably, PC3 delineated an axis separating angiogenesis-related genes with negative loadings (e.g., CAV1, AQP1, CLEC14A) from immune-related genes with positive loadings (e.g., GZMA, CD3E, IR7R), revealing a tumor-associated spatial polarization between vascular remodeling and immune infiltration [47–52]. These genes could be regarded as “spatially differential genes,” highlighting biological processes that are not readily detectable by conventional DEG approaches.

This study had several limitations. First, SCOMV assumes that the reference region forms a sufficiently large mass, which may limit its applicability in cases where the region is highly fragmented into small components or where boundaries with surrounding cell populations are indistinct. Although tumor regions were used as reference regions in this study, SCOMV could be extended to other cell types, such as lymphoid aggregates or neural cell clusters. Second, SCOMV was evaluated using breast and lung cancer datasets, and its generalizability to other tumor types and spatial transcriptomic platforms remain unknown.

Applying this tool to large and diverse datasets may enable *de novo* classification of tumor regions based on spatial gene organization. Integrating tumor-related spatial signatures with clinical annotations could facilitate biomarker discovery and improve our understanding of how tumor architecture is related to disease progression and therapeutic responses.

## Acknowledgements

We thank Dr. Sung Gi Chi, Dr. Yuki Matsubara, Mr. Naoki Amano, Mr. Masaki Ohira, and Ms. Kyoko Kurihara for their valuable comments on this study. We also thank the personnel at the National Cancer Center Hospital East and The University of Tokyo for their assistance and valuable advice.

## Author contributions

Ryosuke Nomura: Conceptualization, Formal analysis, Methodology, Software, Validation, Visualization, Writing – original draft. Shunsuke A. Sakai: Conceptualization, Formal analysis, Writing – review & editing, Shun-Ichiro Kageyama: Conceptualization, Data curation, Funding acquisition, Resources, Validation, Visualization, Writing – review & editing, Katsuya Tsuchihara: Conceptualization, Funding acquisition, Supervision, Validation, Writing – review & editing, Riu Yamashita: Conceptualization, Data curation, Funding acquisition, Project administration, Supervision, Validation, Writing – original draft, Writing – review & editing.

## Data availability

The source code for SpatialCompassV is publicly available at GitHub (https://github.com/RyosukeNomural/SpatialCompassV) and archived in Zenodo under DOI: 10.5281/zenodo.18779150.

## Appendix

Not applicable.

