## Supplementary Figures S1-S39 for "SpatialCompassV (SCOMV): *De novo* cell and gene spatial pattern classification and spatially differential gene identification"

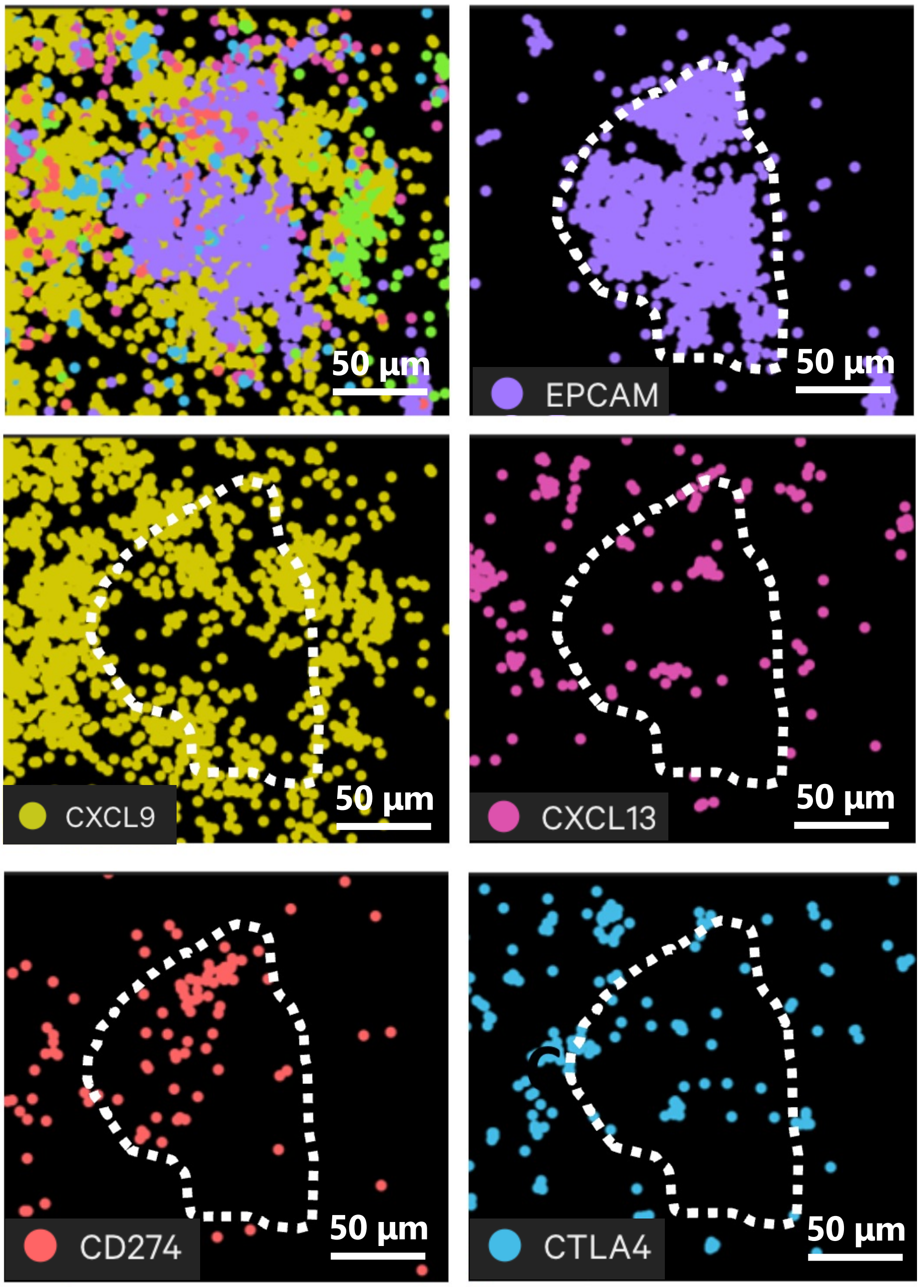


**Supplementary Fig. S1 Visualization of selected genes in Xenium Explore.**
Spatial expression patterns of EPCAM, CXCL9, CXCL13, CD274 and CTLA4 displayed using Xenium Explore. White lines indicate the tumor boundary.

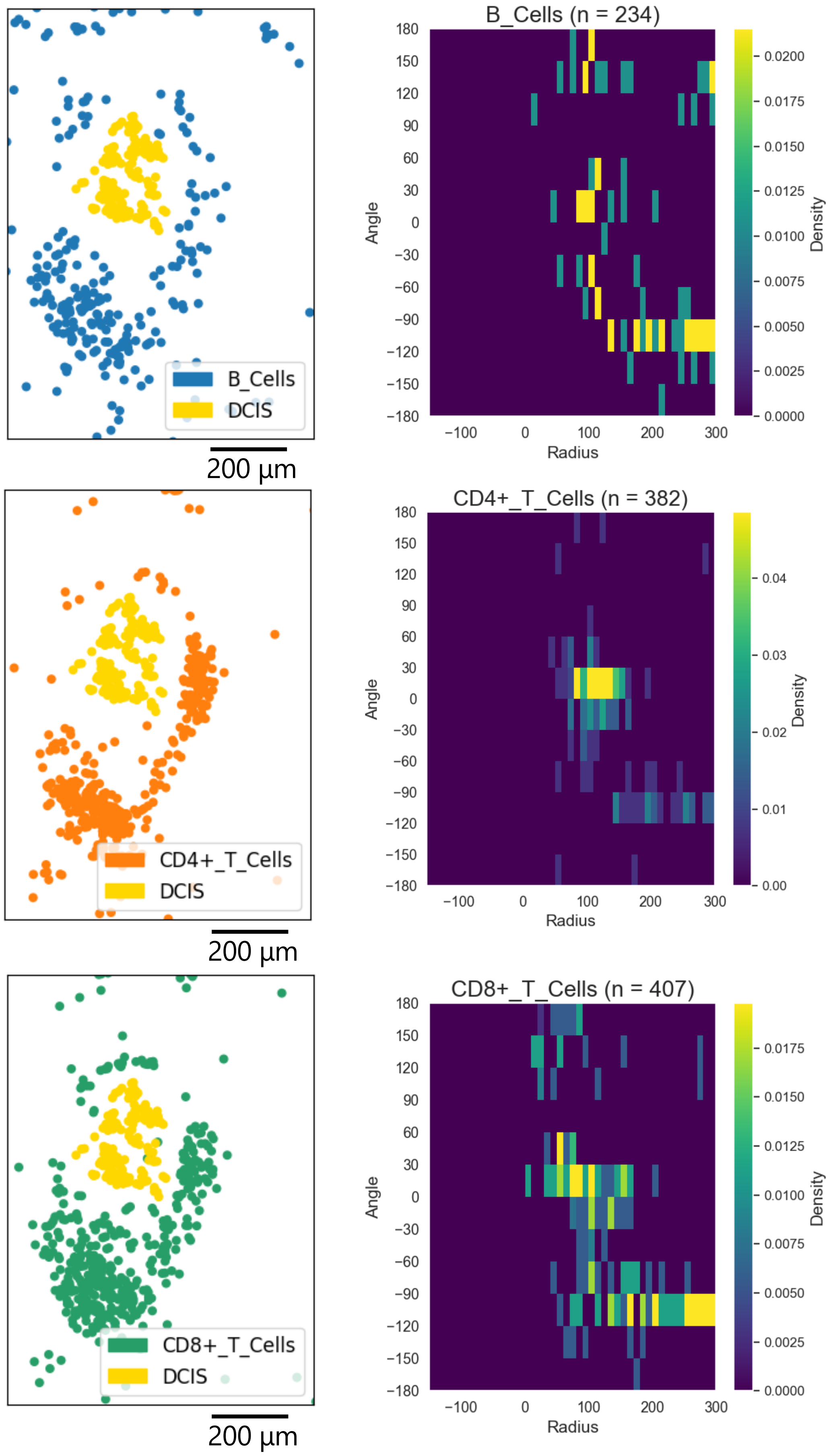


**Supplementary Fig. S2 Visualization of selected cells and feature matrix.**
Spatial expression patterns of B cells, $CD4^+$ T cells, and $CD8^+$ T cells were visualized using Python (Scanpy). The feature matrix was plotted using Python (matplotlib); the x-axis shows the distance from the tumor region (negative values indicate intratumoral positions), and the y-axis shows the angle.

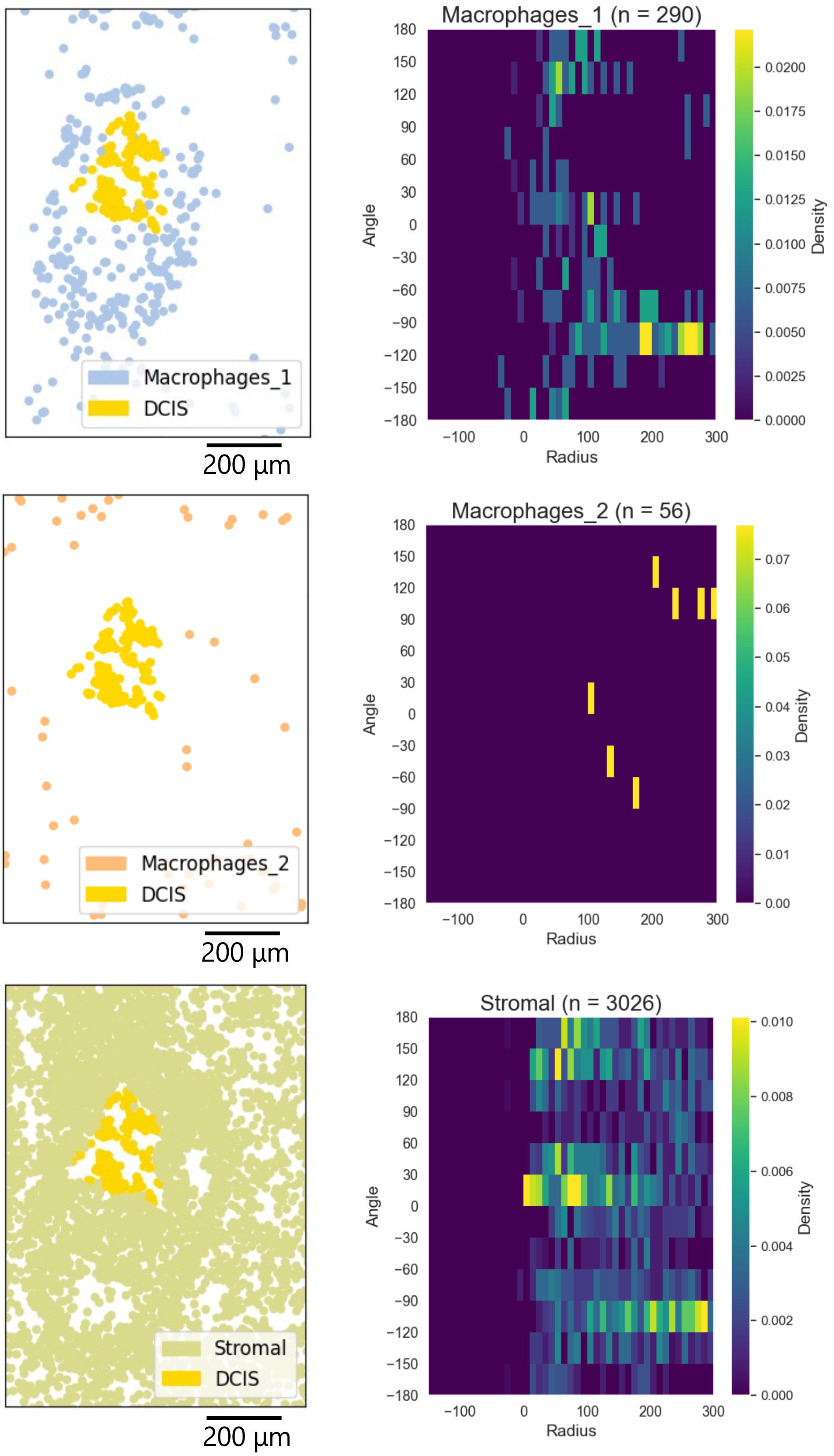


**Supplementary Fig. S3 Visualization of selected cells and feature matrix.**
Spatial expression patterns of M1 macrophages, $CD4^{+}$, M2 macrophages and Stromal cells were visualized using Python (Scanpy). The feature matrix was plotted using Python (matplotlib); the x-axis shows the distance from the tumor region (negative values indicate intratumoral positions), and the y-axis shows the angle.

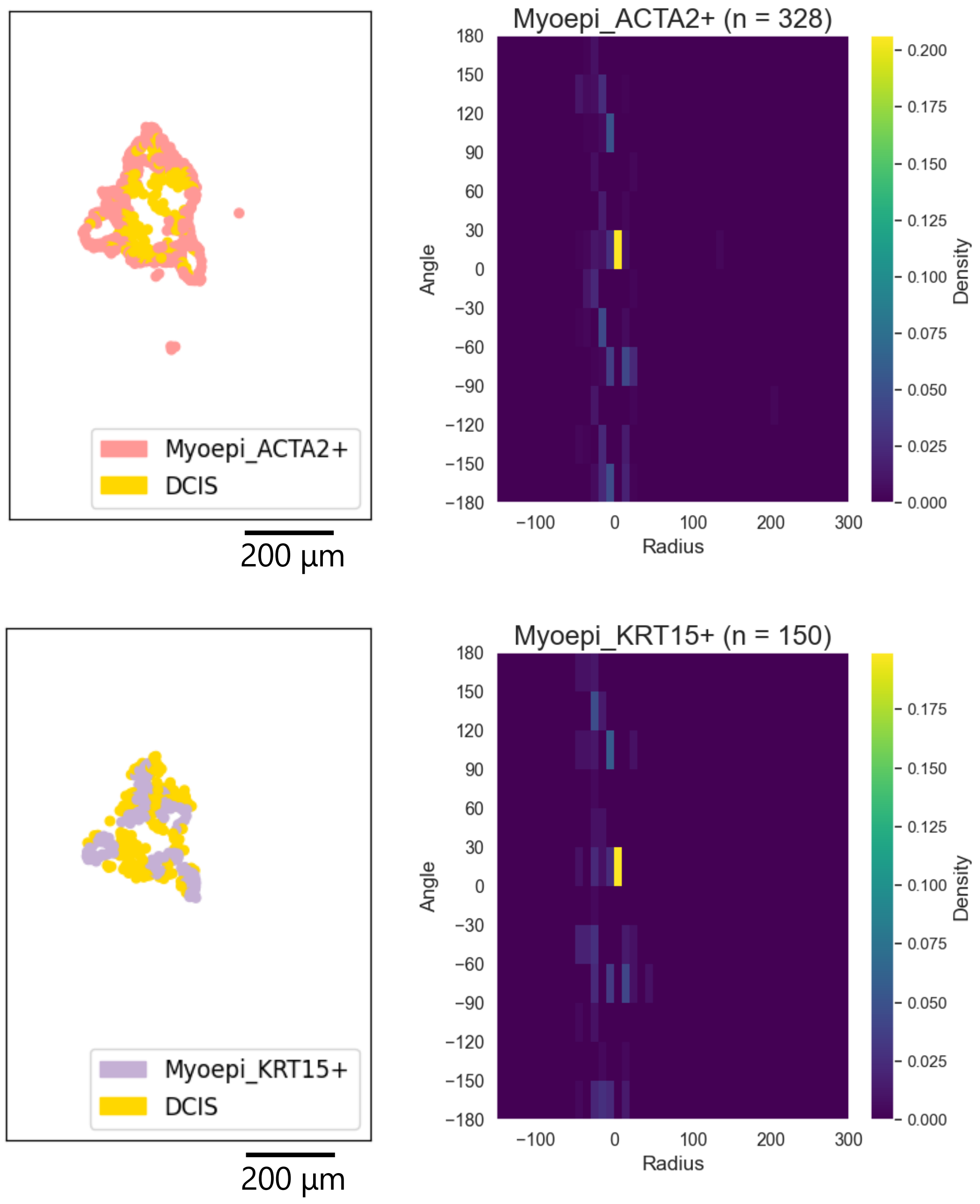


**Supplementary Fig. S4 Visualization of selected cells and feature matrix.**

Spatial expression patterns of Myoepithelial ACTA2$^+$ and Myoepithelial KRT15$^+$ cells were visualized using Python (Scanpy). The feature matrix was plotted using Python (matplotlib); the x-axis shows the distance from the tumor region (negative values indicate intratumoral positions), and the y-axis shows the angle.

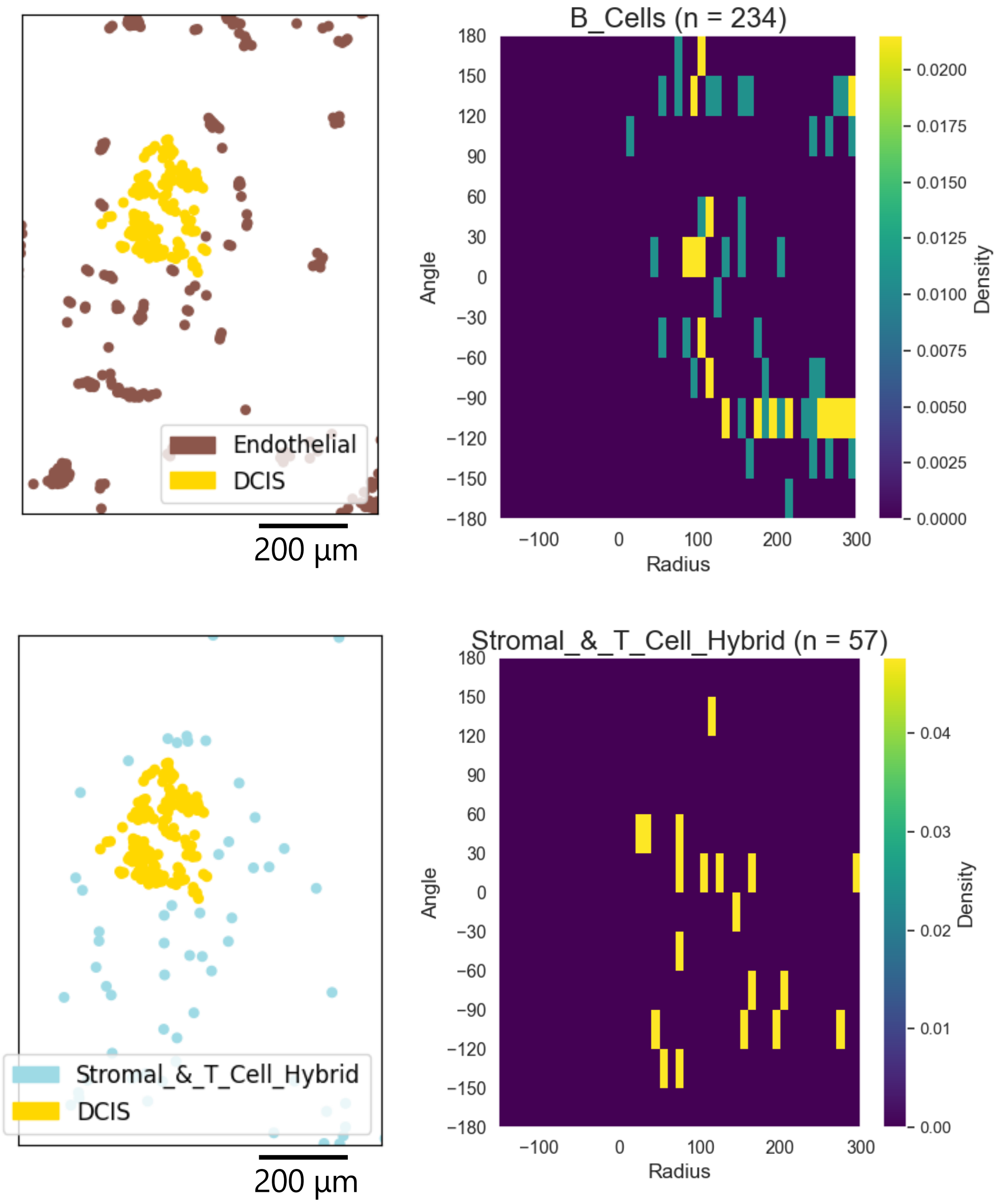


**Supplementary Fig. S5 Visualization of selected cells and feature matrix.**
Spatial expression patterns of Endothelial and stromal/T-cell hybrid cells were visualized using Python (Scanpy). The feature matrix was plotted using Python (matplotlib); the x-axis shows the distance from the tumor region (negative values indicate intratumoral positions), and the y-axis shows the angle.

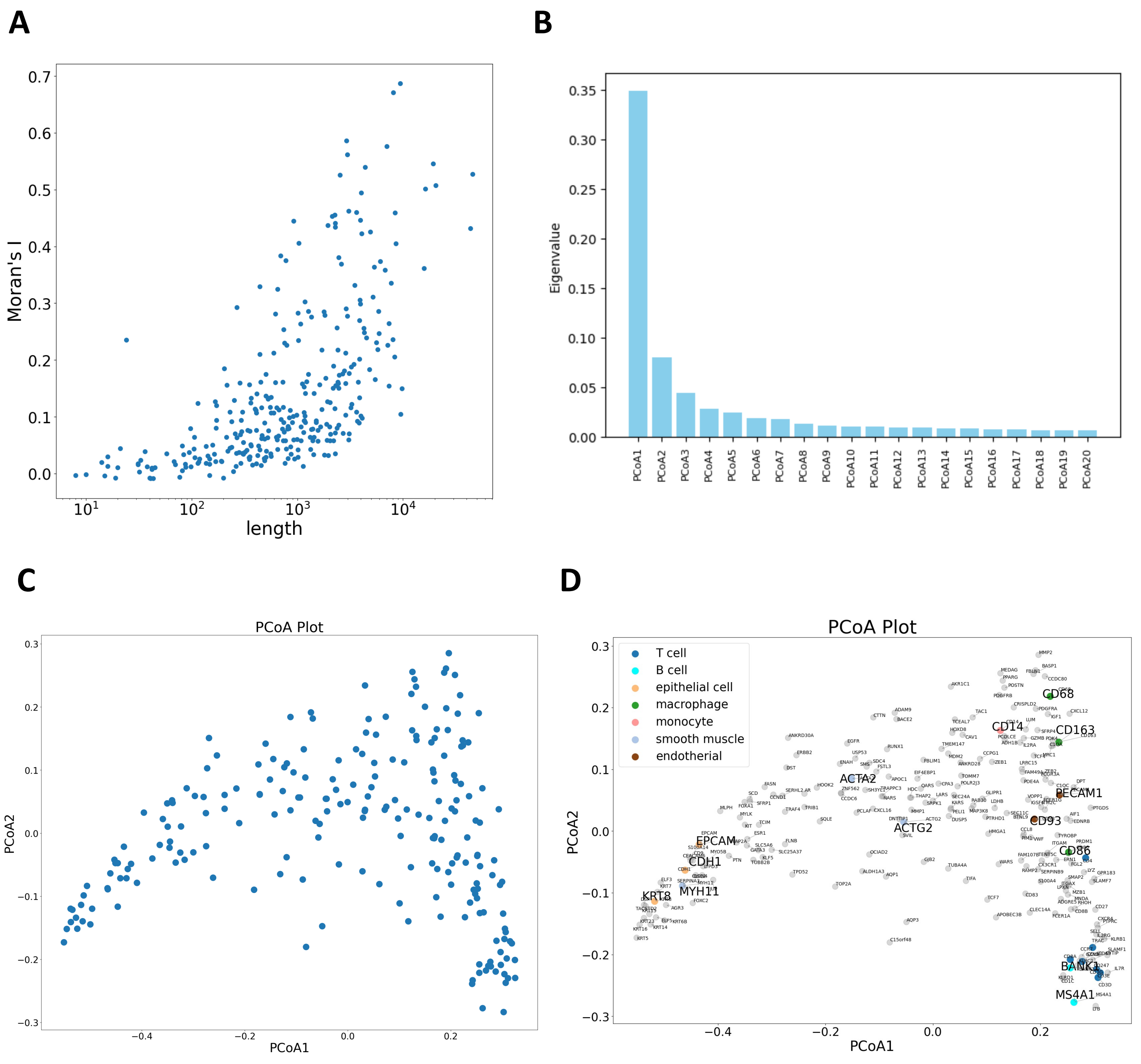


**Supplementary Fig. S6 Detailed results of gene–gene similarity in the breast cancer panel.** (A) Relationship between gene expression levels and Moran's I. (B) Principal coordinate analysis (PCoA) of the gene-gene similarity matrix. Axes represent PCoA components with the proportion of variance explained. (C) Projection of the first two PCoA components. (D) Gene name annotation of the PCoA plot in (C). Uppercase labels correspond to those in Fig. 3A.

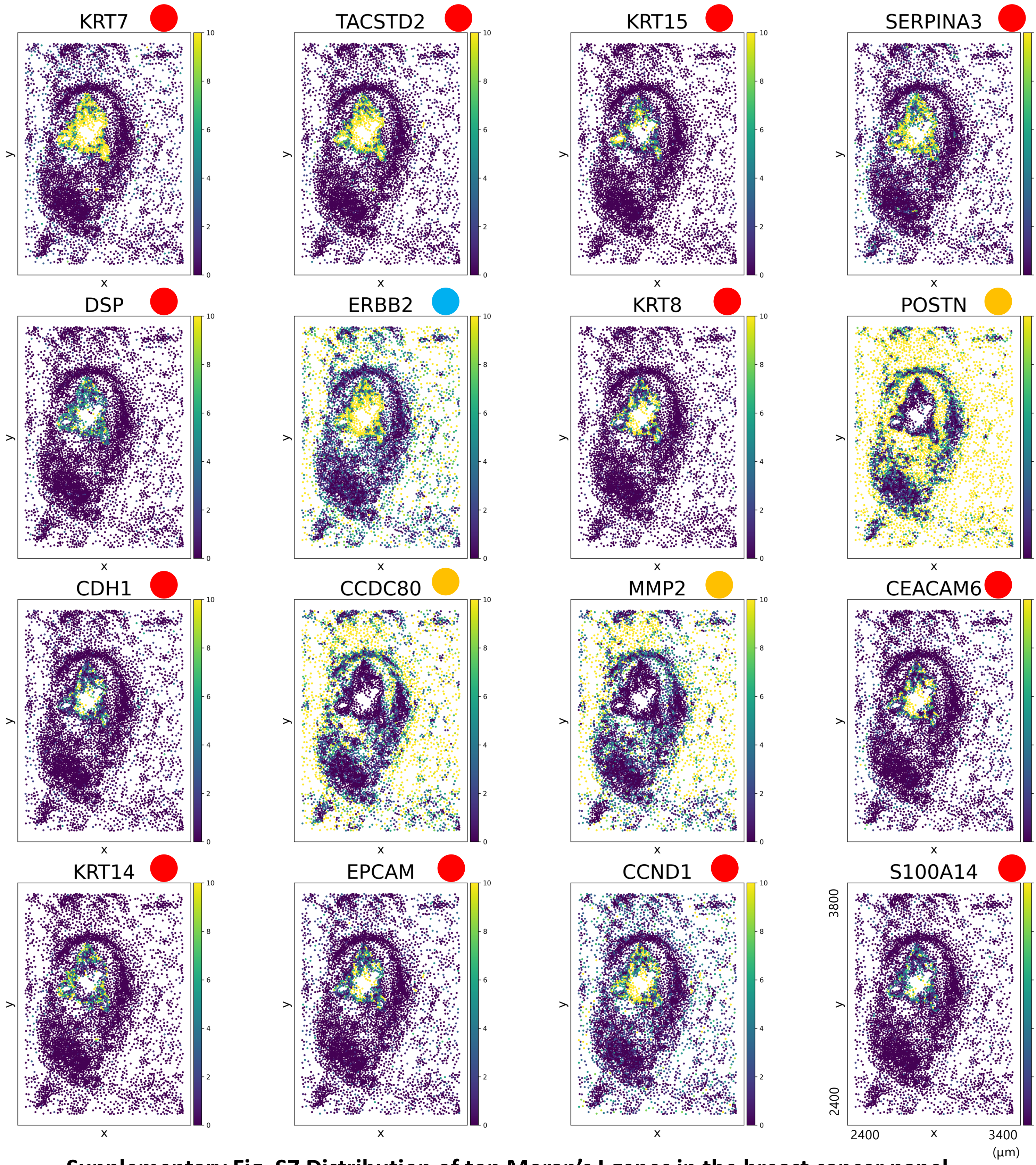


**Supplementary Fig. S7 Distribution of top Moran's I genes in the breast cancer panel.** Spatial expression maps for the top 16 genes ranked by Moran's I in the breast cancer panel were generated using Python (Squidpy) and arranged from the upper left to the lower right. Circles next to gene names indicate spatial patterns (red: internal, orange: peripheral, blue: ubiquitous). Moran's I values do not necessarily concord with tumor-relative spatial distributions.

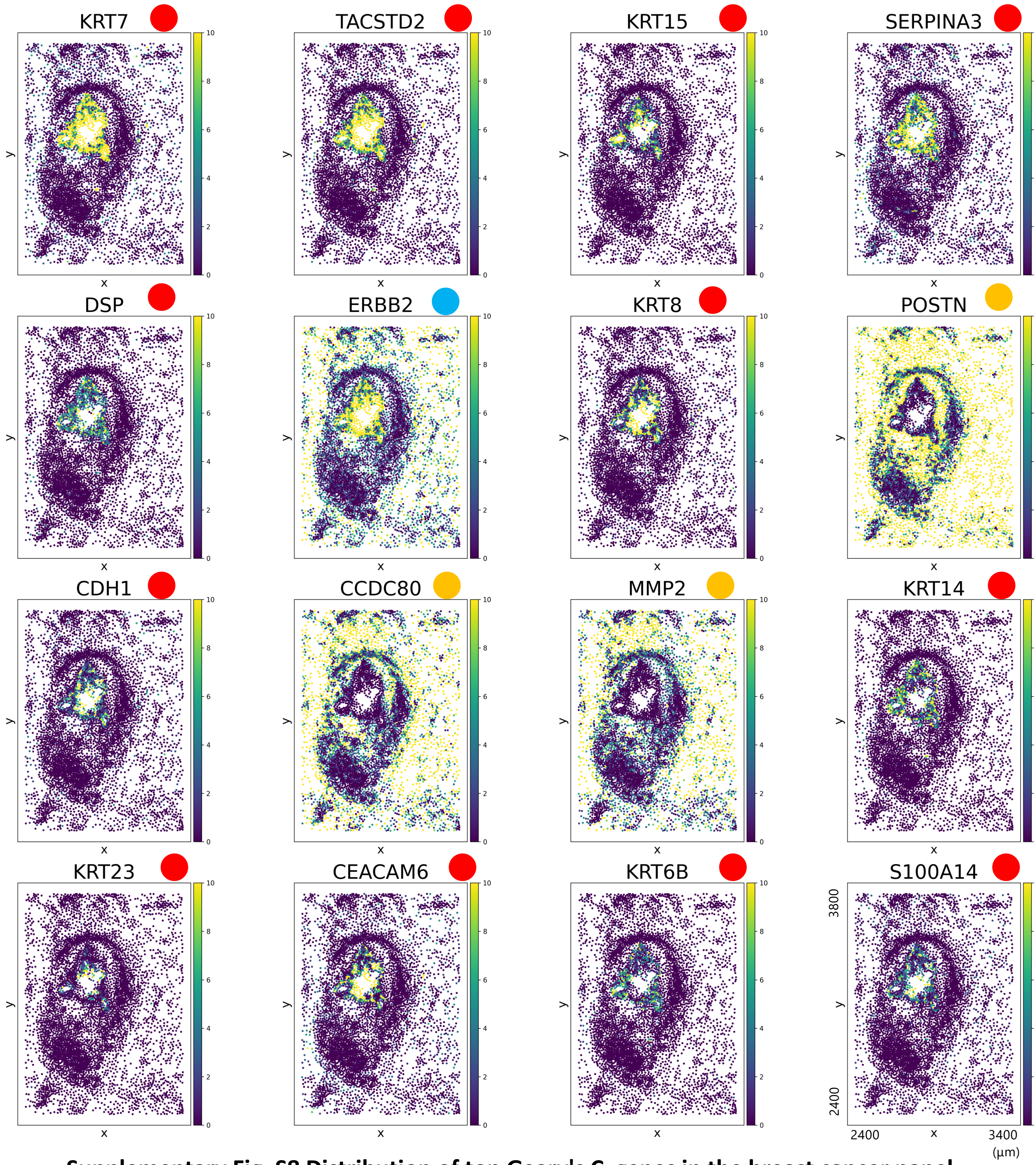


**Supplementary Fig. S8 Distribution of top Geary's C genes in the breast cancer panel.** Spatial expression maps for the top 16 genes ranked by Geary's C in the breast cancer panel were generated using Python (Squidpy) and arranged from the upper left to the lower right. Circles next to gene names indicate spatial patterns (red: internal, orange: peripheral, blue: ubiquitous). Geary's C values do not necessarily concord with tumor-relative spatial distributions.

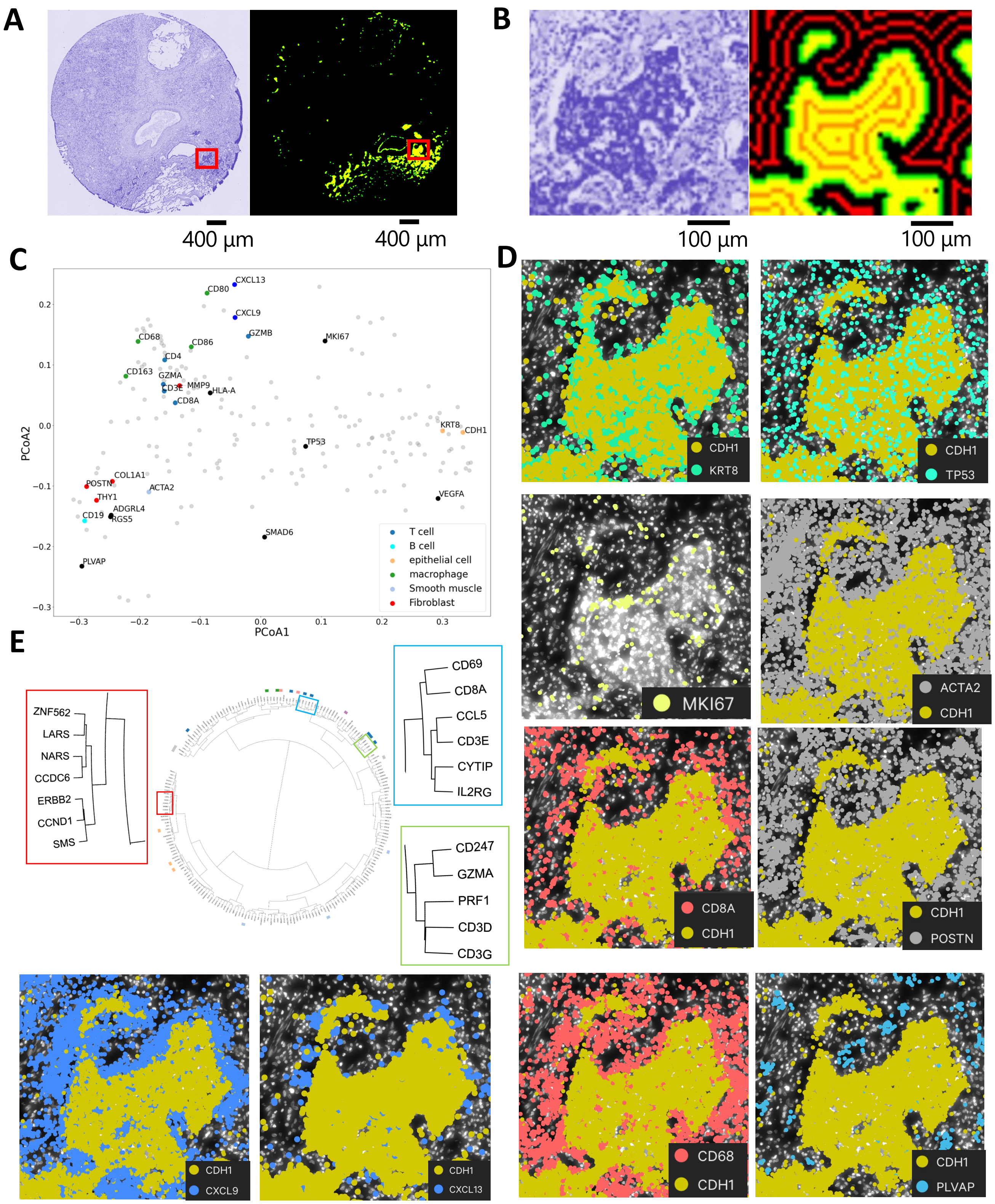


**Supplementary Fig. S9 Gene distribution analysis in lung cancer panel.**
(A) Hematoxylin and eosin (HE) image (left) and tumor regions extracted by SpatialKnifeY(right), with the analyzed region of interest (ROI, red box). (B) Enlarged view of the selected ROI. (C) PCoA based on the similarity matrix of gene distribution patterns, where each point represents a gene. (D) Spatial expression patterns of representative genes selected from PCoA space. CDH1 is used to indicate the epithelial tumor region. (E) Hierarchical clustering of the gene–gene similarity matrix shown as a dendrogram. The Newick-formatted tree was visualized using iTOL. Boxed regions indicate magnified views. Blue highlights denote immune cell markers (e.g. CD8A, CD3E), yellow indicates cytotoxic genes (e.g. PRF1, GZMA) and red marks functionally related genes (e.g. LARS, NARS) clustered in close proximity.

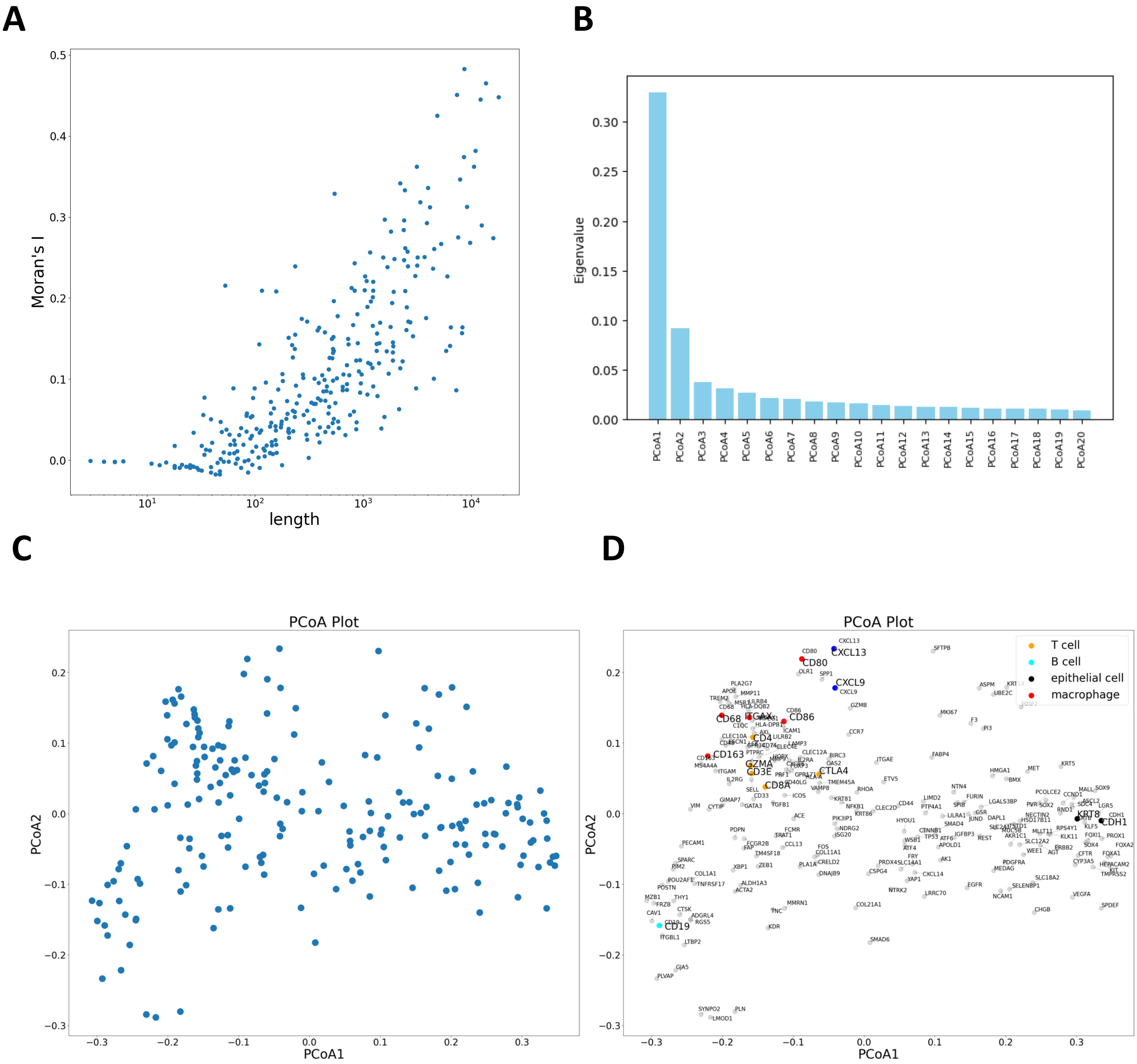


**Supplementary Fig. S1- Detailed results of gene–gene similarity in the lung cancer panel.** (A) Relationship between gene expression levels and Moran's I. (B) Principal coordinate analysis (PCoA) of the gene-gene similarity matrix. Axes represent PCoA components with the proportion of variance explained. (C) Projection of the first two PCoA components. (D) Gene name annotation of the PCoA plot in (C).

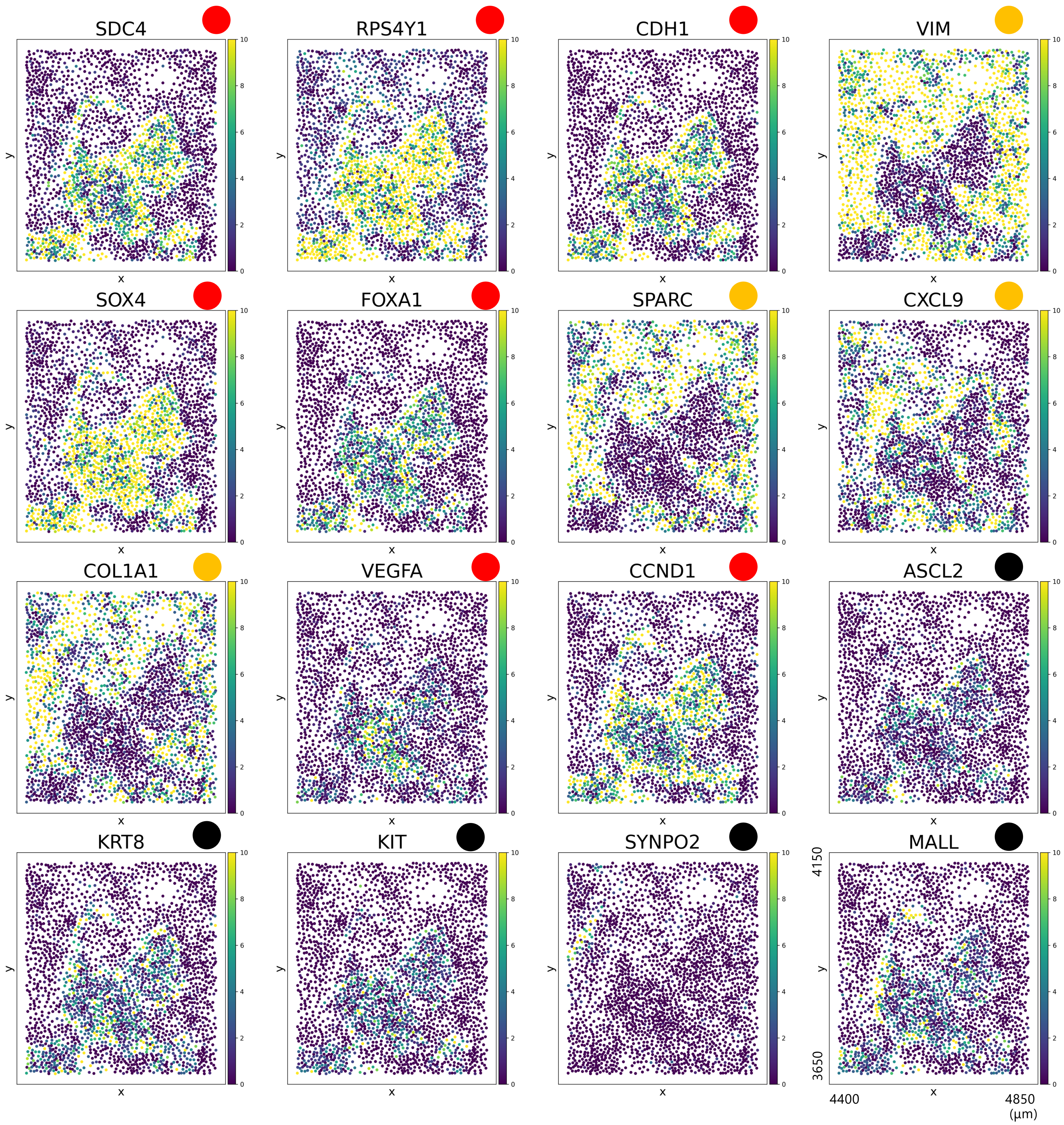


**Supplementary Fig. S11 Distribution of top Moran's I genes in the breast cancer panel.** Spatial expression maps for the top 16 genes ranked by Moran's I in the breast cancer panel were generated using Python (Squidpy) and arranged from the upper left to the lower right. Circles next to gene names indicate spatial patterns (red: internal, orange: peripheral, black: low expression). Moran's I values do not necessarily concord with tumor-relative spatial distributions.

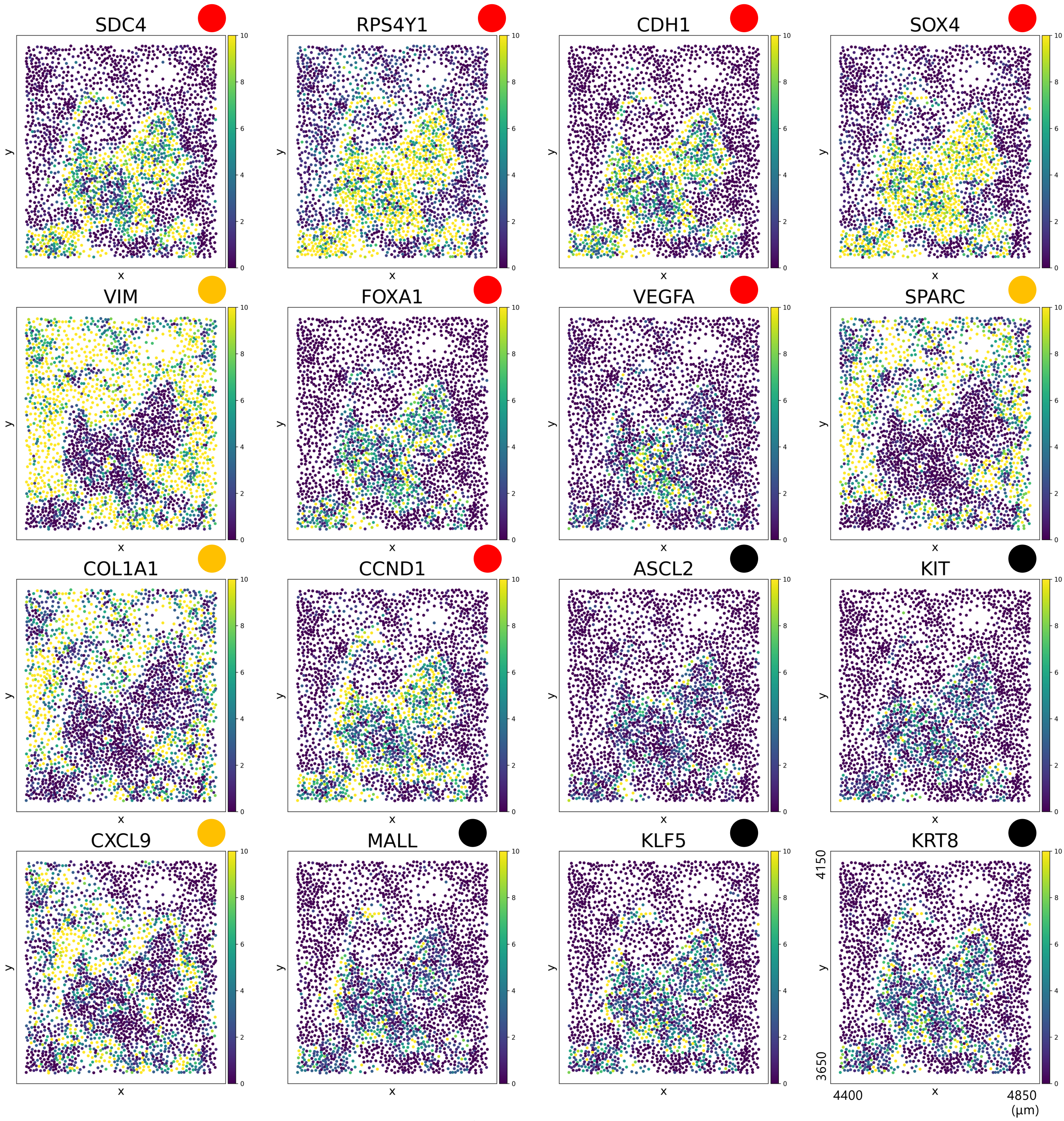


**Supplementary Fig. S12 Distribution of top Geary's C genes in the breast cancer panel.** Spatial expression maps for the top 16 genes ranked by Geary's C in the breast cancer panel were generated using Python (Squidpy) and arranged from the upper left to the lower right. Circles next to gene names indicate spatial patterns (red: internal, orange: peripheral, black: low expression). Geary's C values do not necessarily concord with tumor-relative spatial distributions.

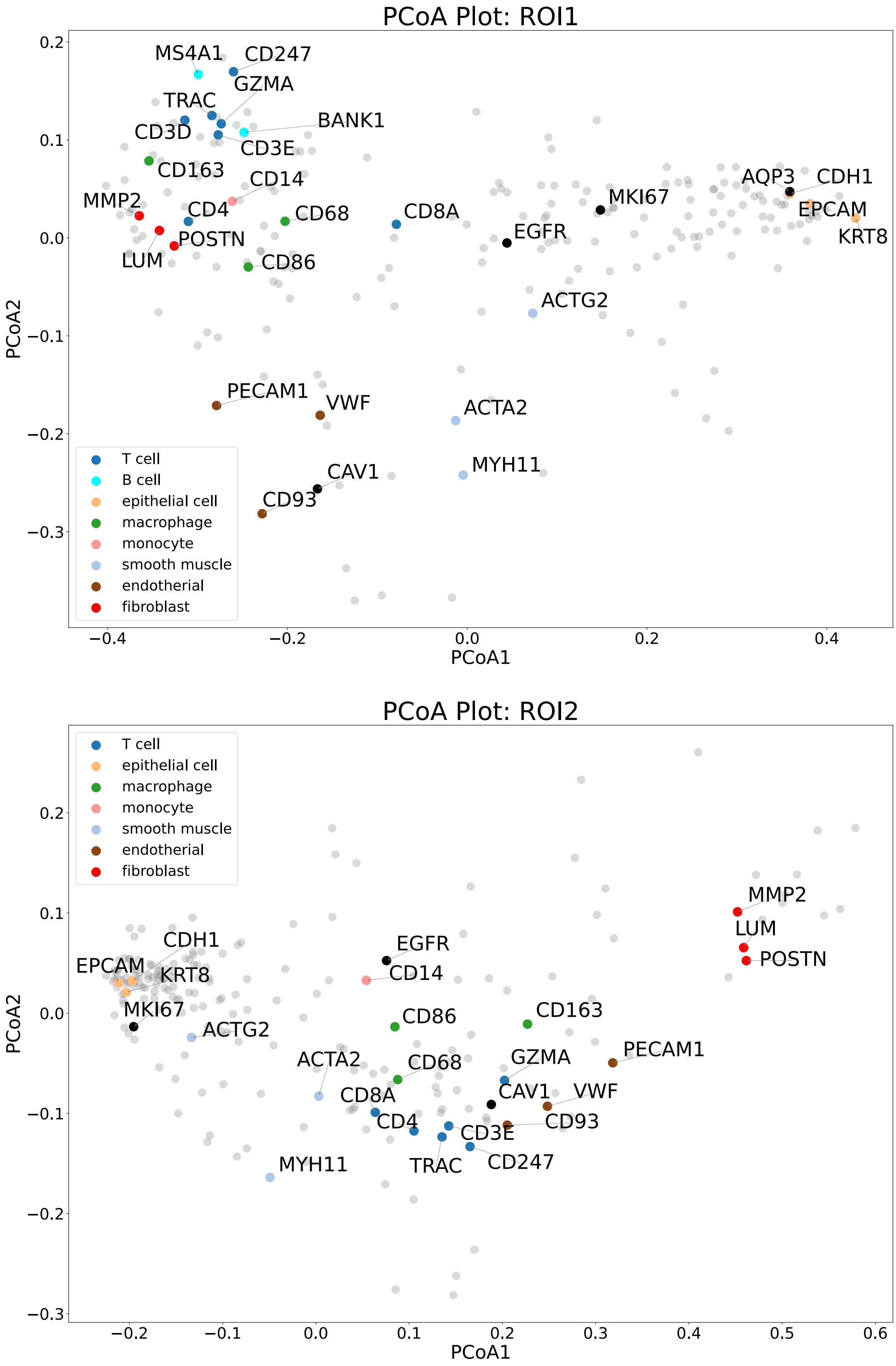


**Supplementary Fig. S13 PCoA of ROI1 and ROI2 in the multiple-field analysis.** Two-dimensional PCoA plots of ROI1 and ROI2 used in the multiple-field analysis in Fig. 4. Points are colored according to cell type–specific marker gene sets.

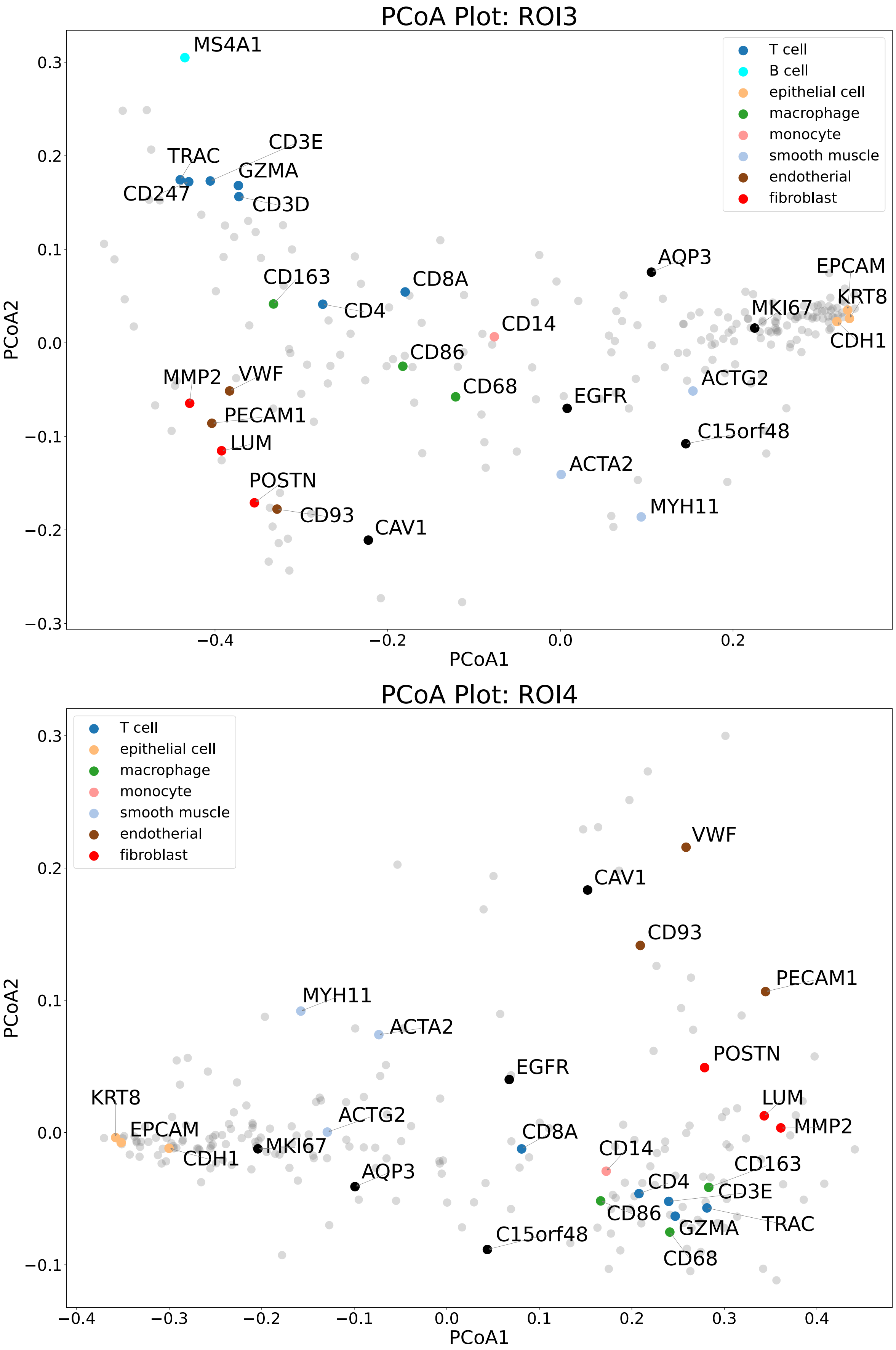


**Supplementary Fig. S14 PCoA of ROI3 and ROI4 in the multiple-field analysis.** Two-dimensional PCoA plots of ROI3 and ROI4 used in the multiple-field analysis in Fig. 4. Points are colored according to cell type–specific marker gene sets.

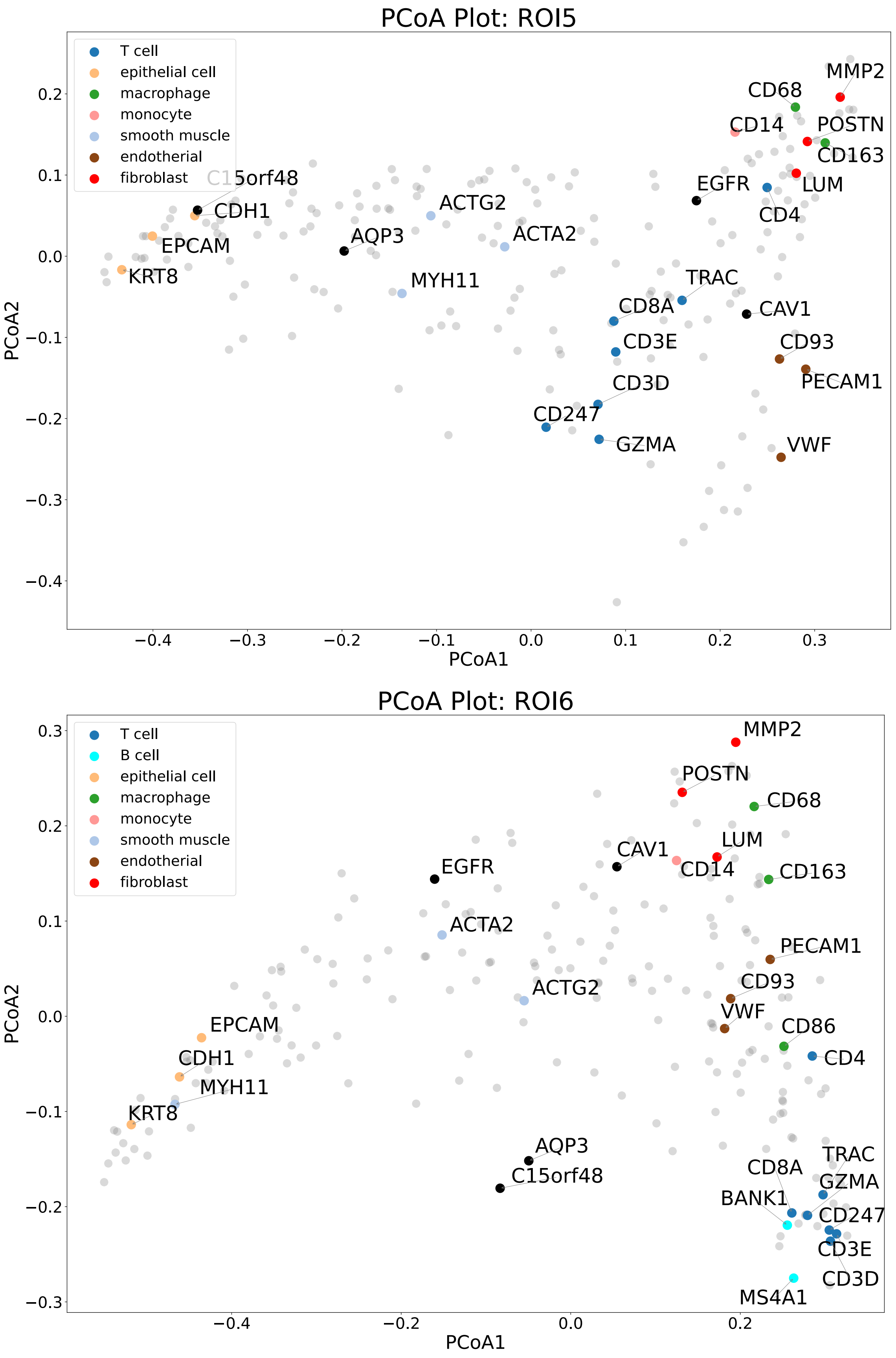


**Supplementary Fig. S15 PCoA of ROI5 and ROI6 in the multiple-field analysis.** Two-dimensional PCoA plots of ROI5 and ROI6 used in the multiple-field analysis in Fig. 4. Points are colored according to cell type–specific marker gene sets.

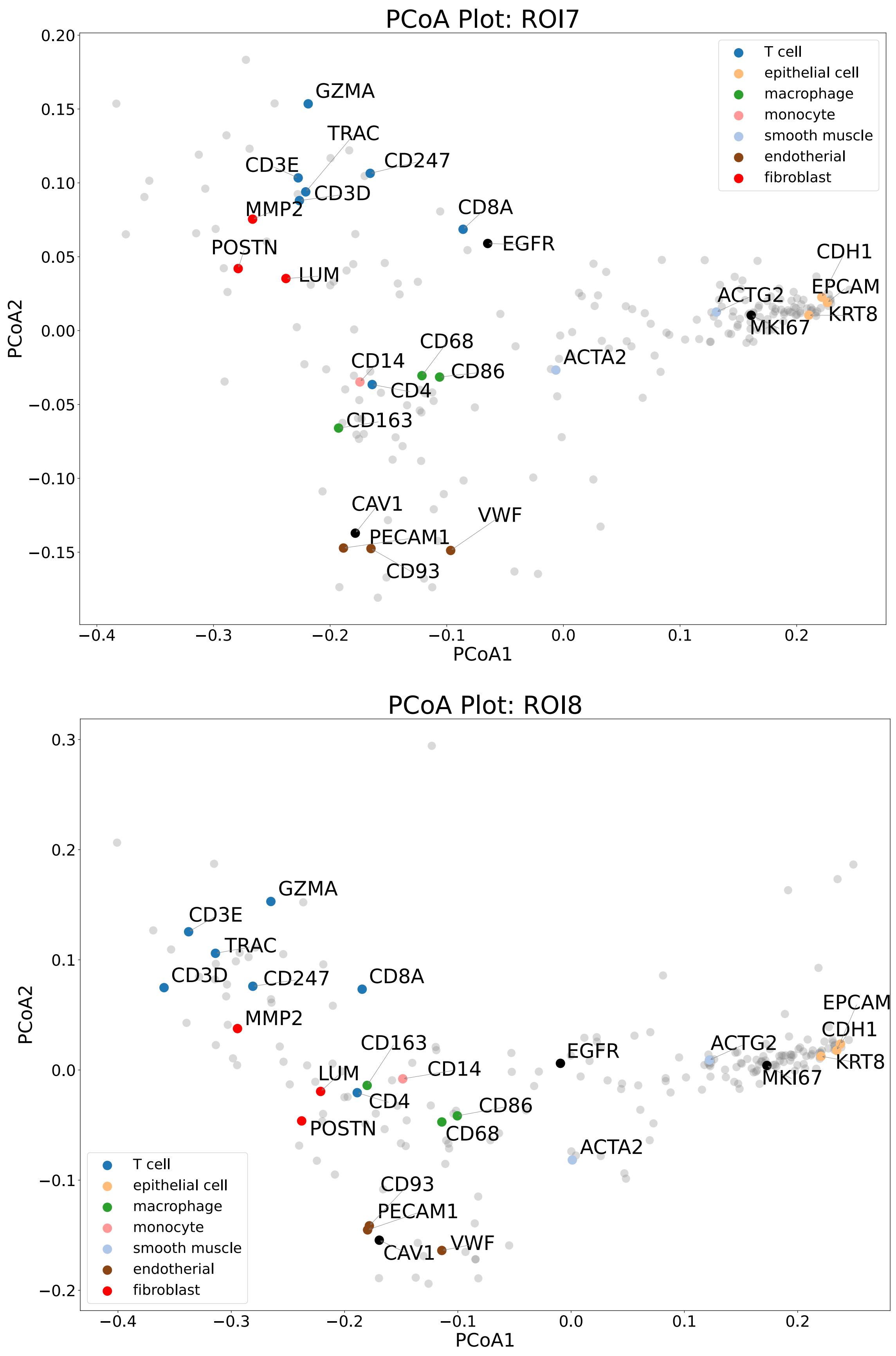


**Supplementary Fig. S16 PCoA of ROI7 and ROI8 in the multiple-field analysis.** Two-dimensional PCoA plots of ROI7 and ROI8 used in the multiple-field analysis in Fig. 4. Points are colored according to cell type–specific marker gene sets.

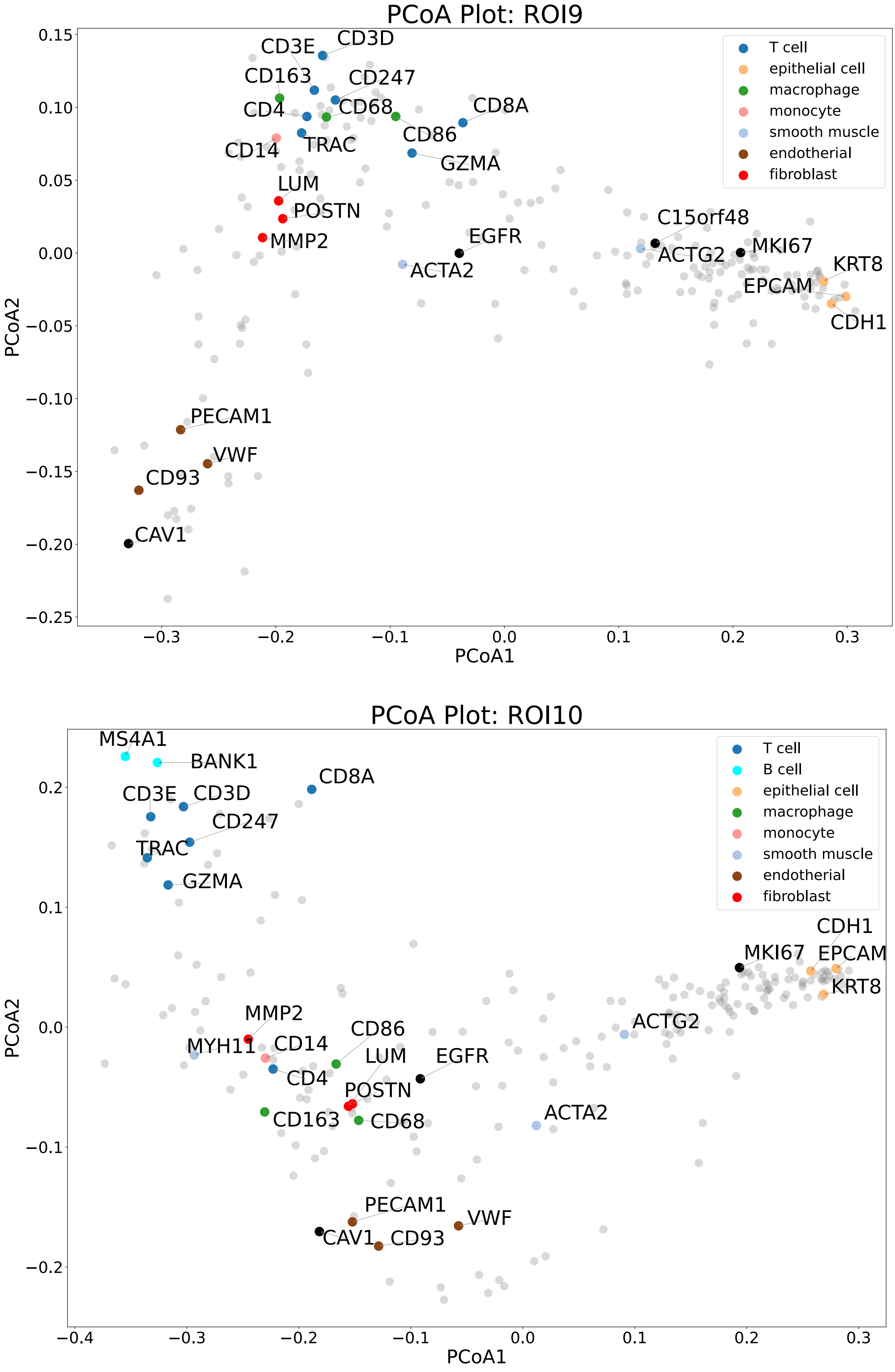


**Supplementary Fig. S17 PCoA of ROI9 and ROI10 in the multiple-field analysis.** Two-dimensional PCoA plots of ROI9 and ROI10 used in the multiple-field analysis in Fig. 4. Points are colored according to cell type–specific marker gene sets.

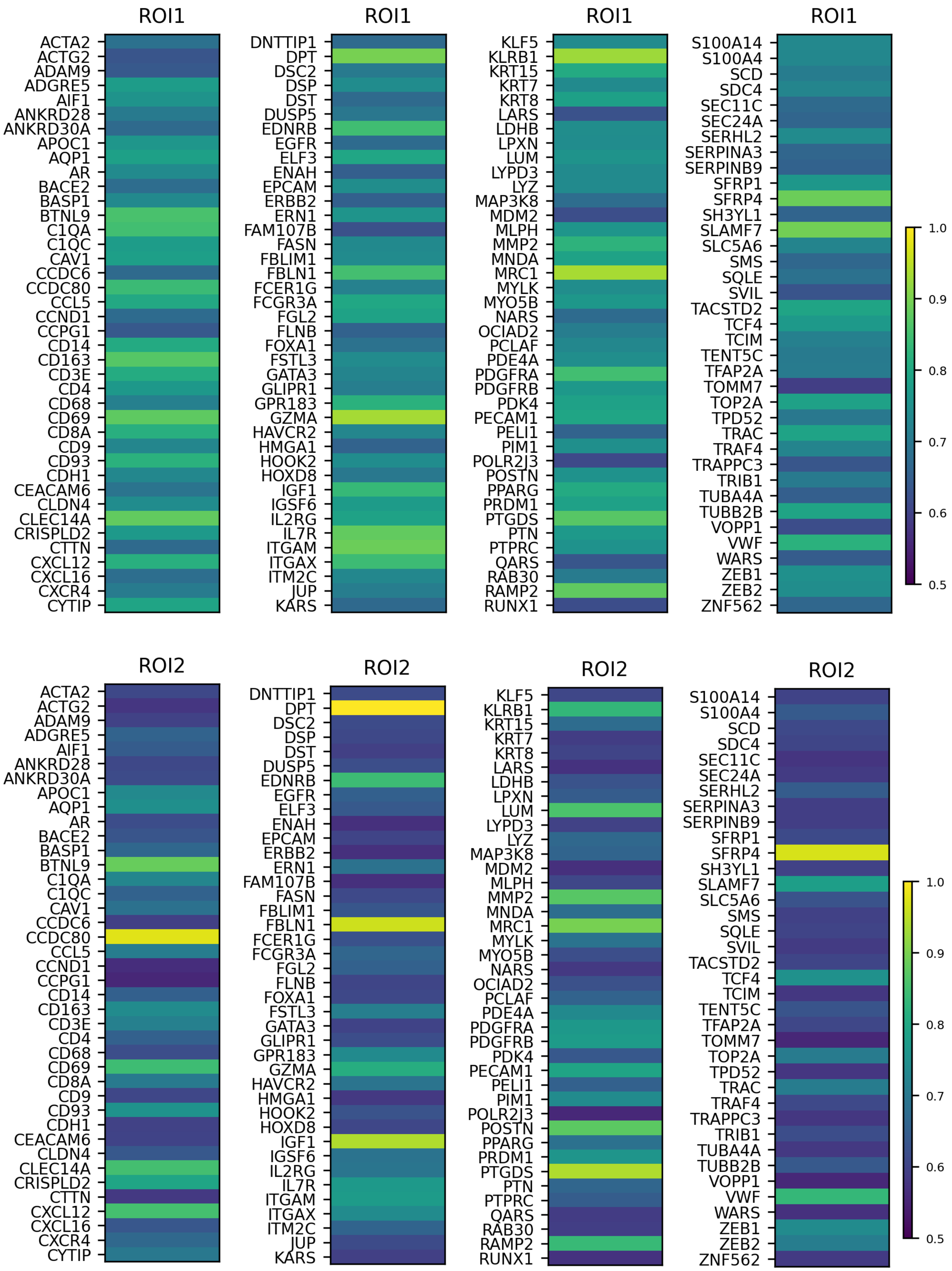


**Supplementary Fig. S18 One-dimensional symmetric NMF representation of gene–gene similarity in ROI1 and ROI2.**

Gene–gene similarity matrices from ROI1 and ROI2 were reduced to one-dimensional vectors by symmetric non-negative matrix factorization (NMF) and plotted using Python (matplotlib). Gene names are shown on the y-axis in alphabetical order.

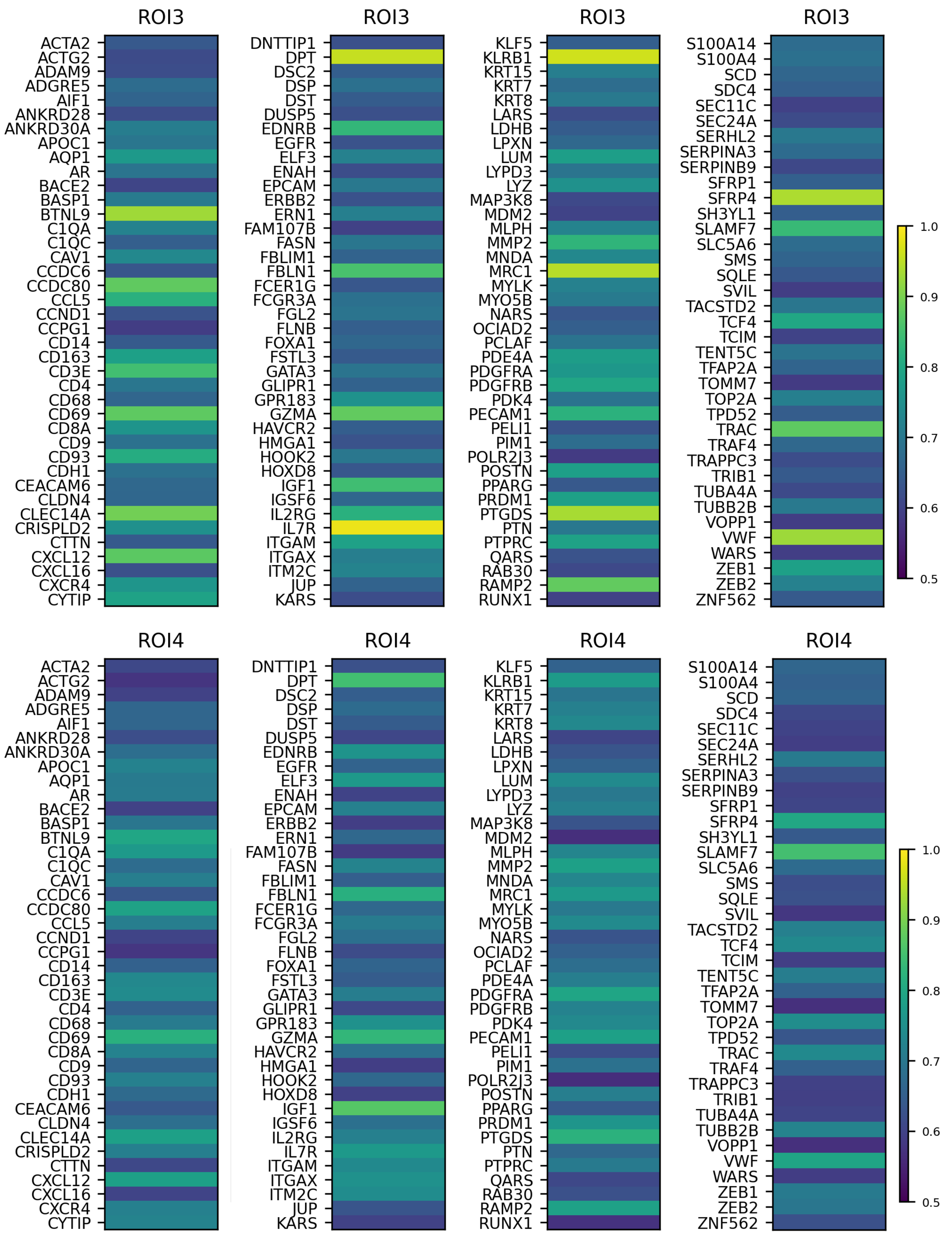


**Supplementary Fig. S19 One-dimensional symmetric NMF representation of gene–gene similarity in ROI3 and ROI4.**

Gene–gene similarity matrices from ROI3 and ROI4 were reduced to one-dimensional vectors by symmetric non-negative matrix factorization (NMF) and plotted using Python (matplotlib). Gene names are shown on the y-axis in alphabetical order.

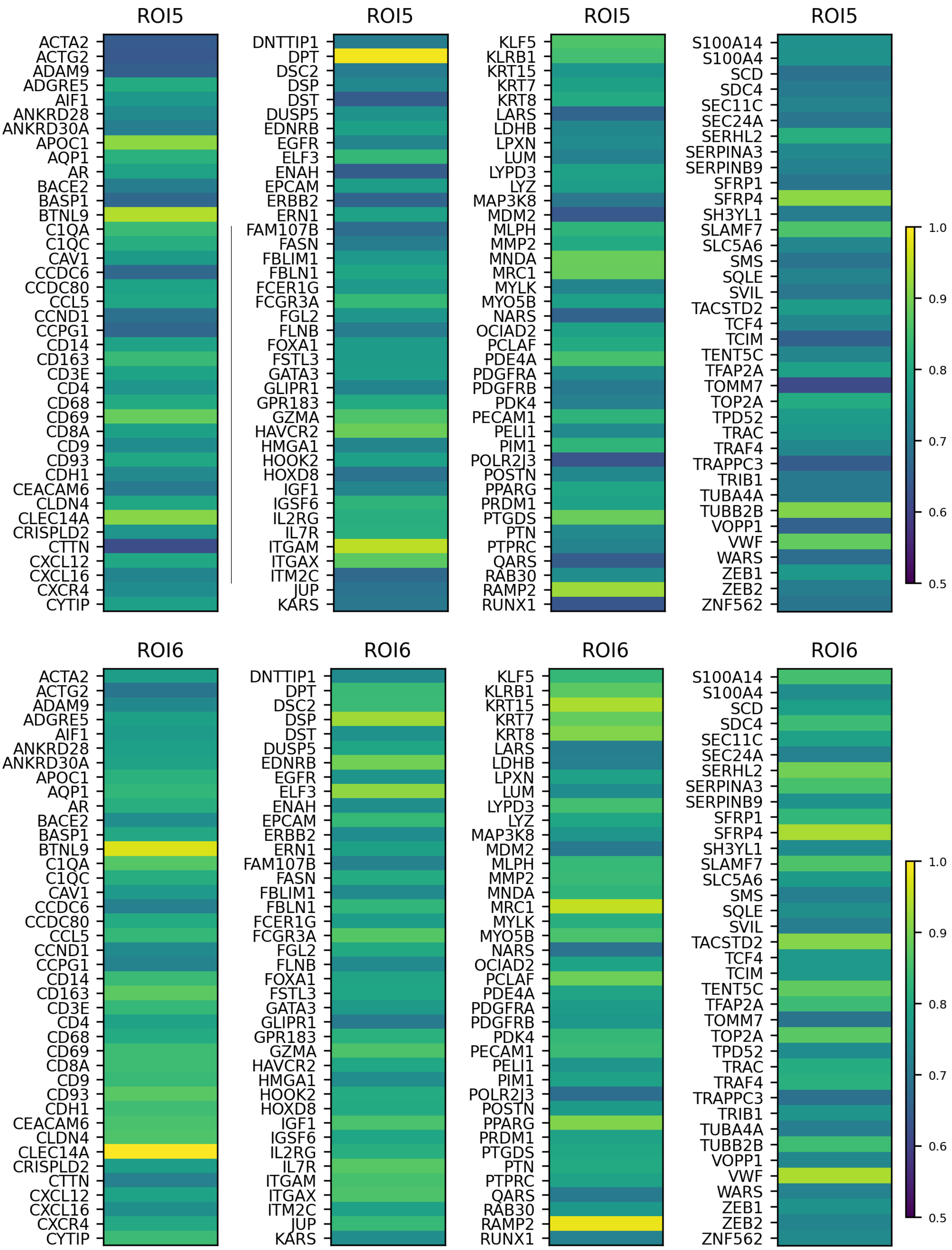


**Supplementary Fig. S20 One-dimensional symmetric NMF representation of gene–gene similarity in ROI5 and ROI6.**

Gene–gene similarity matrices from ROI5 and ROI6 were reduced to one-dimensional vectors by symmetric non-negative matrix factorization (NMF) and plotted using Python (matplotlib). Gene names are shown on the y-axis in alphabetical order.

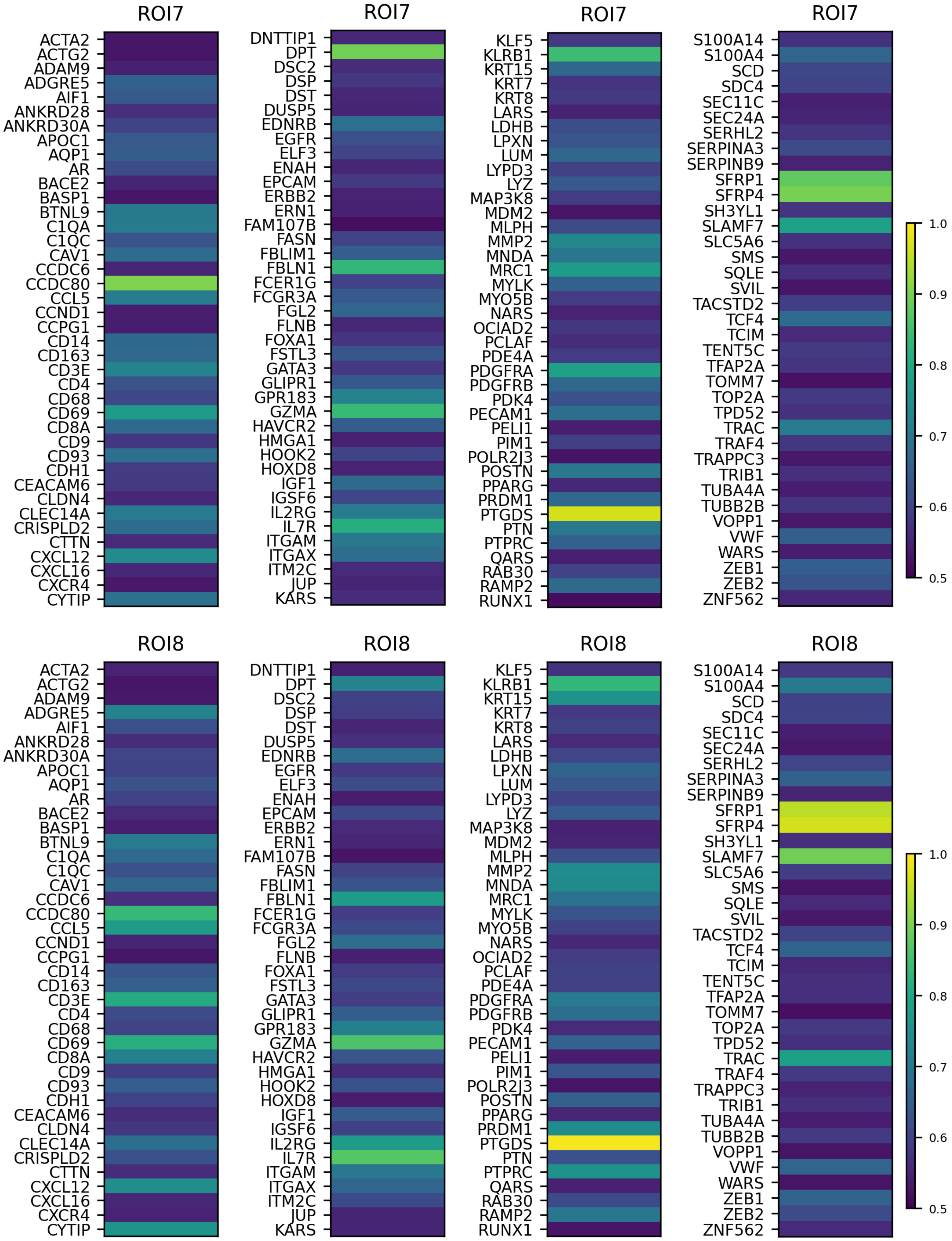


**Supplementary Fig. S21 One-dimensional symmetric NMF representation of gene–gene similarity in ROI7 and ROI8.**

Gene–gene similarity matrices from ROI7 and ROI8 were reduced to one-dimensional vectors by symmetric non-negative matrix factorization (NMF) and plotted using Python (matplotlib). Gene names are shown on the y-axis in alphabetical order.

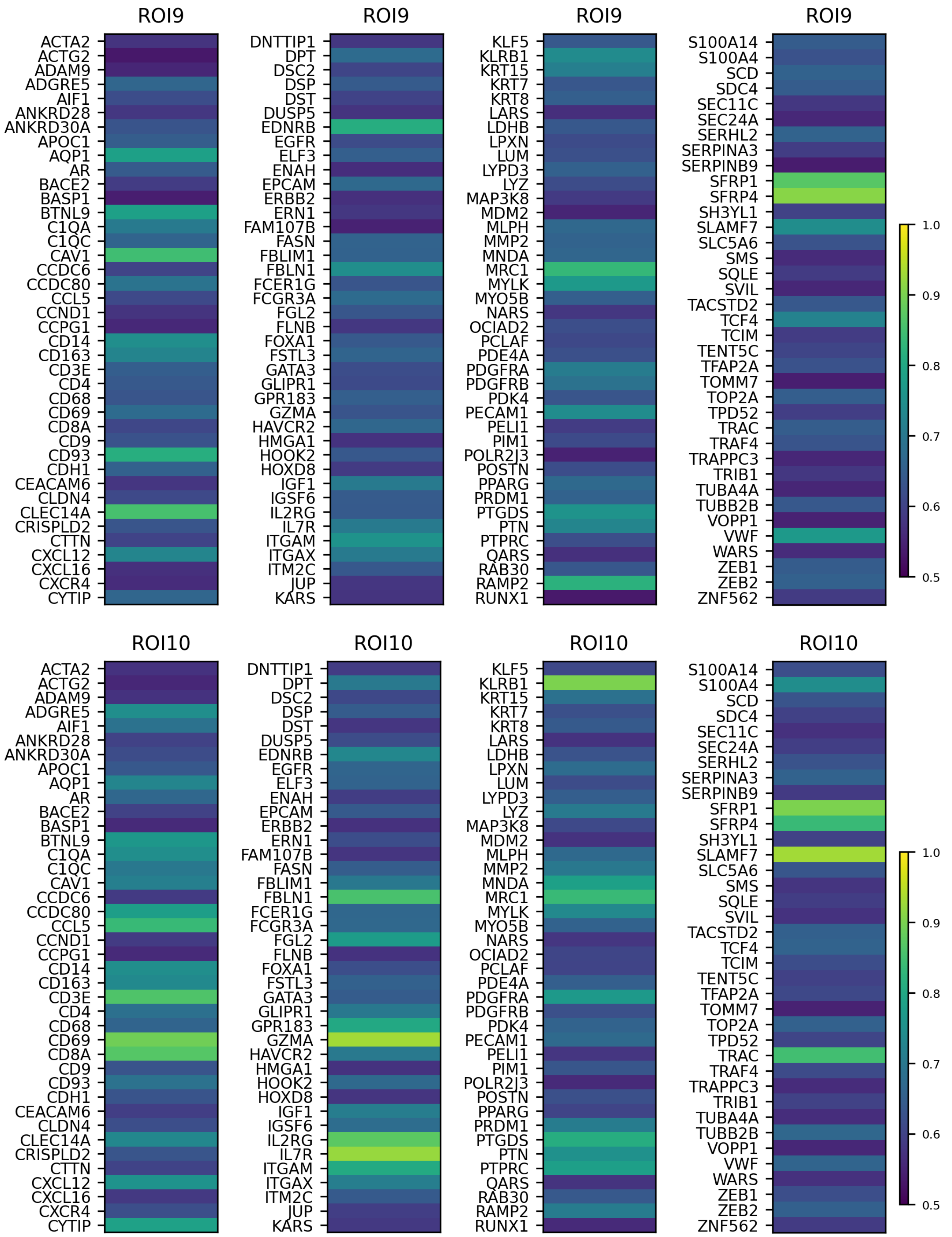


**Supplementary Fig. S22 One-dimensional symmetric NMF representation of gene–gene similarity in ROI9 and ROI10.**

Gene–gene similarity matrices from ROI8 and ROI10 were reduced to one-dimensional vectors by symmetric non-negative matrix factorization (NMF) and plotted using Python (matplotlib). Gene names are shown on the y-axis in alphabetical order.

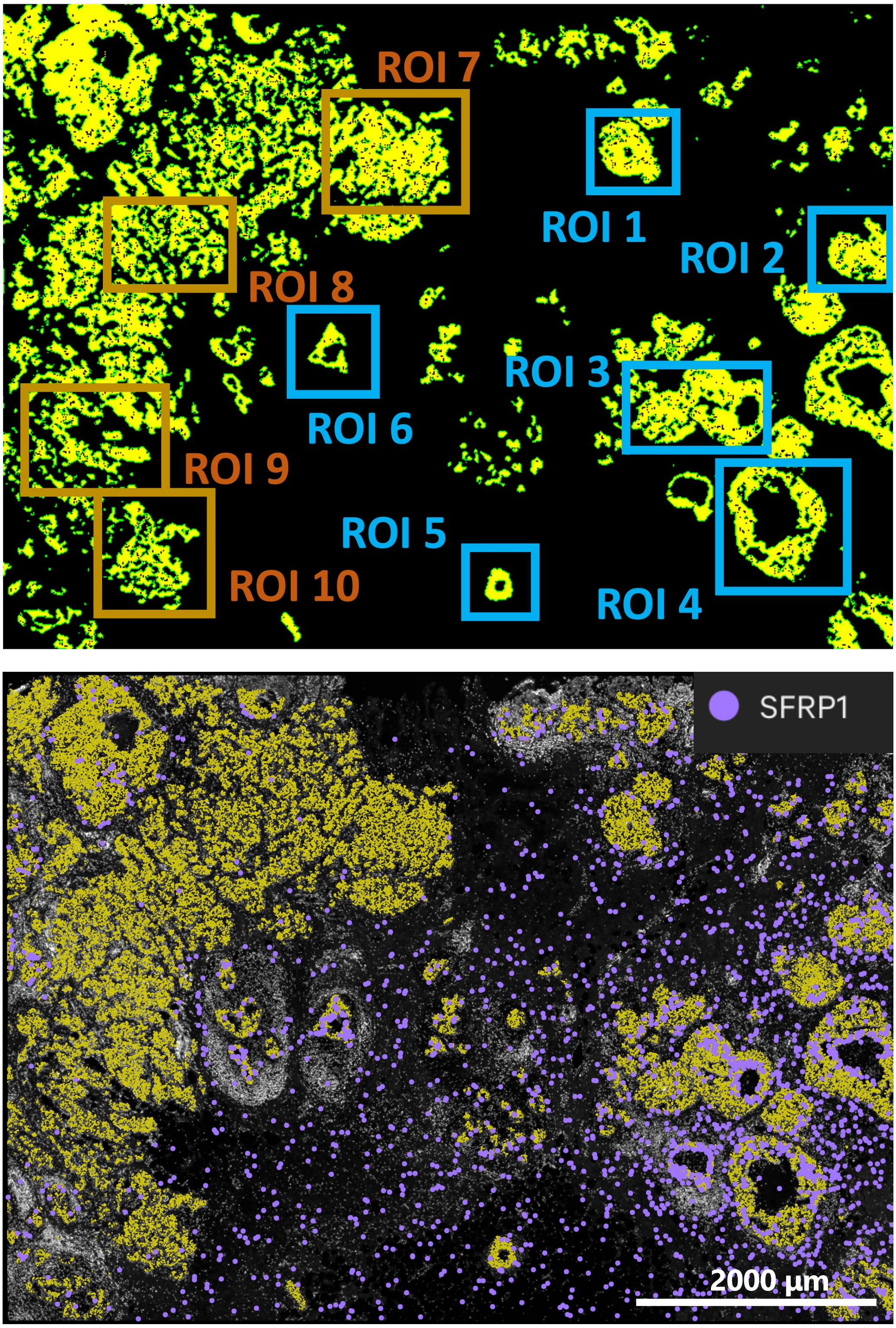


**Supplementary Fig. S23 Selected ROIs and spatial distribution of SFRP1.**
The upper panels show tumor regions identified by SpatialKnifeY (yellow) and the selected regions of interest (ROIs). The lower panels show the spatial expression of SFRP1, visualized using Xenium Explorer.

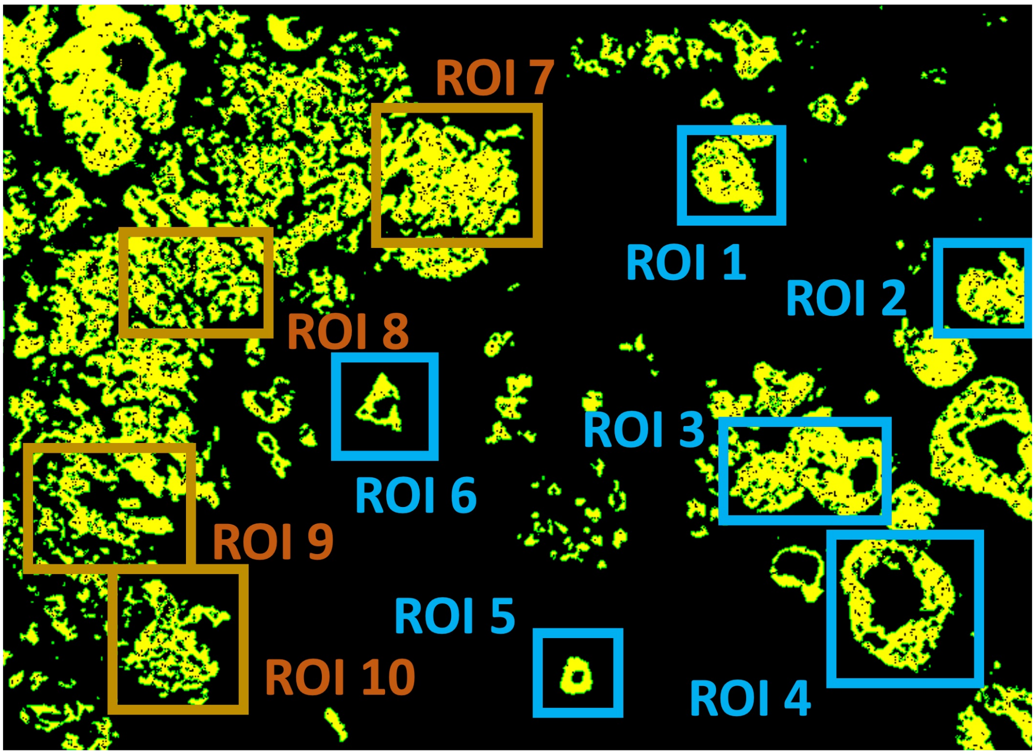


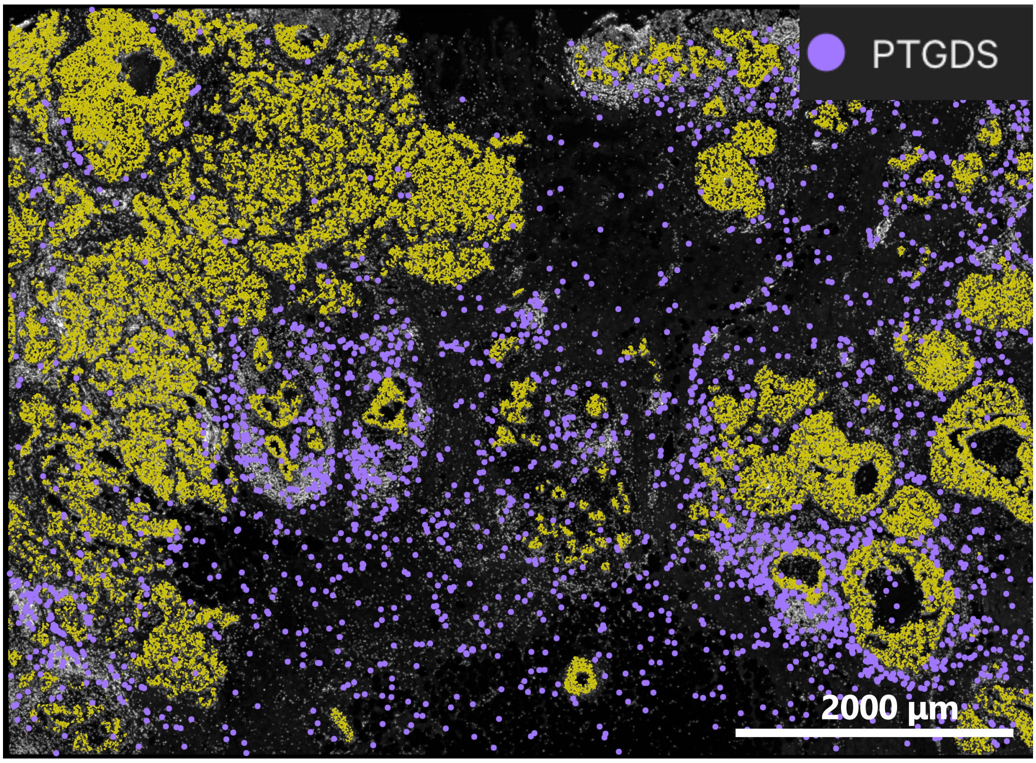


**Supplementary Fig. S24 Selected ROIs and spatial distribution of PTGDS.**
The upper panels show tumor regions identified by SpatialKnifeY (yellow) and the selected regions of interest (ROIs). The lower panels show the spatial expression of PTGDS, visualized using Xenium Explorer.

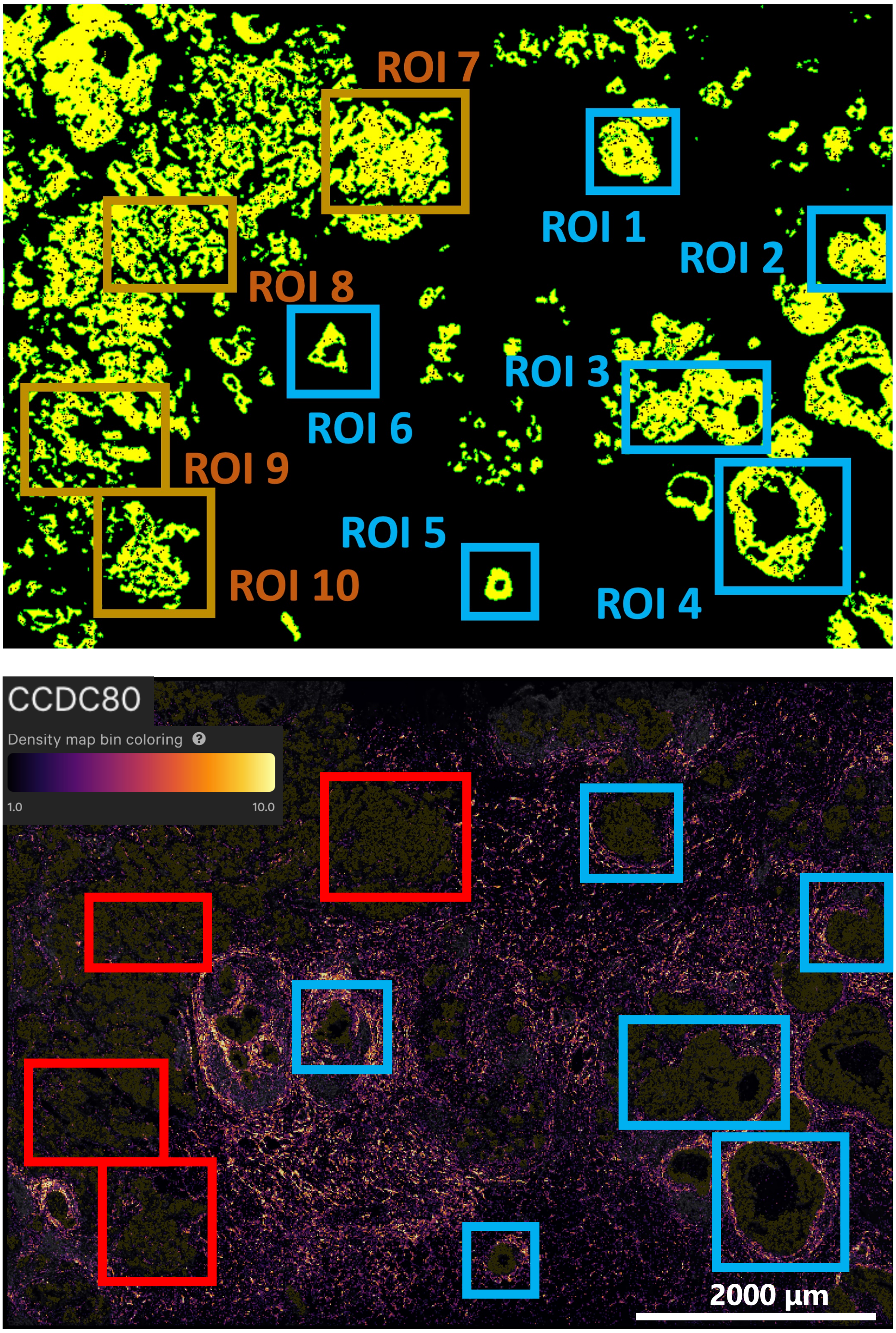


**Supplementary Fig. S25 Selected ROIs and spatial distribution of CCDC80.**
The upper panels show tumor regions identified by SpatialKnifeY (yellow) and the selected regions of interest (ROIs). The lower panels show the spatial expression of CCDC80, visualized using Xenium Explorer.

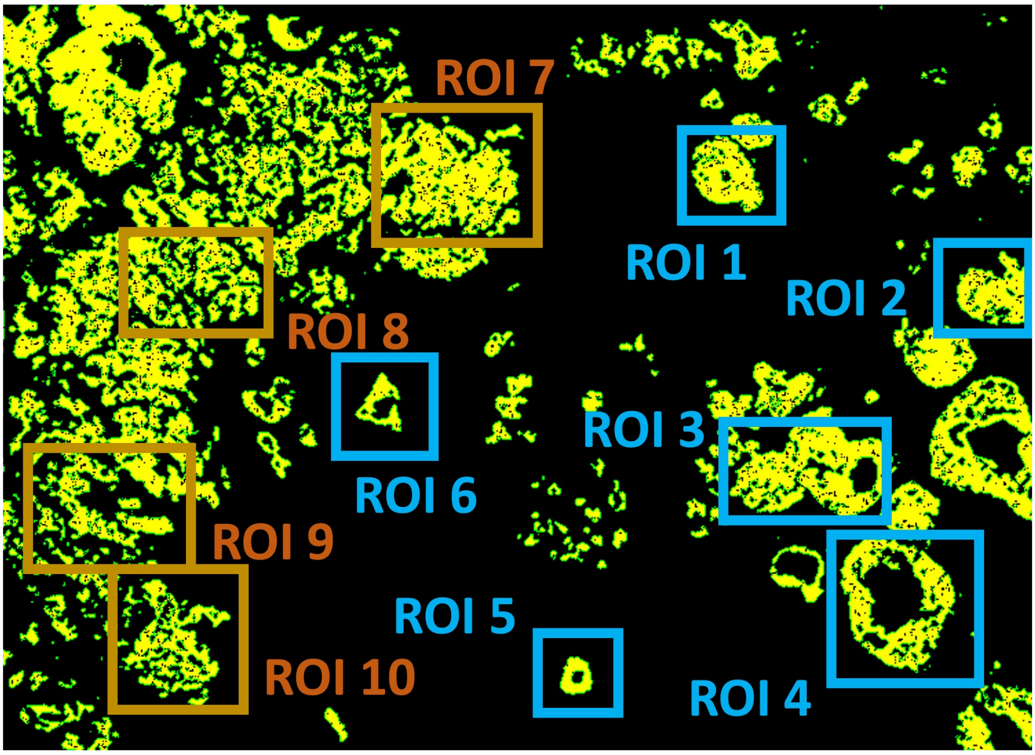


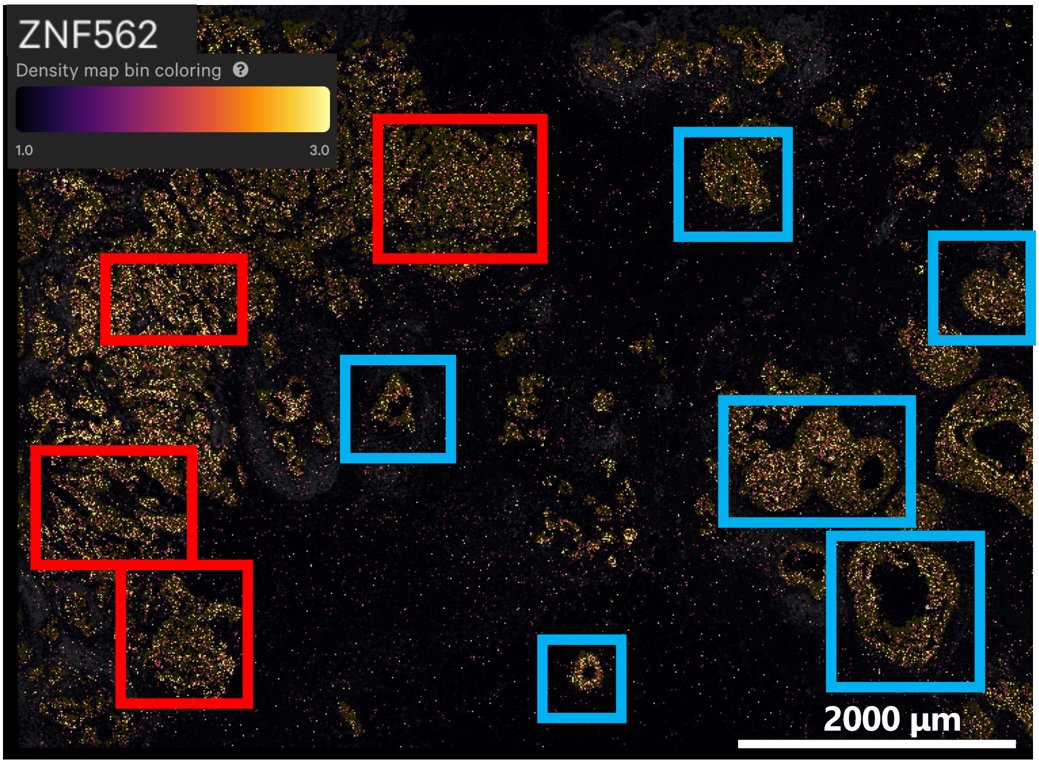


**Supplementary Fig. S26 Selected ROIs and spatial distribution of ZNF562.**
The upper panels show tumor regions identified by SpatialKnifeY (yellow) and the selected regions of interest (ROIs). The lower panels show the spatial expression of ZNF562, visualized using Xenium Explorer.

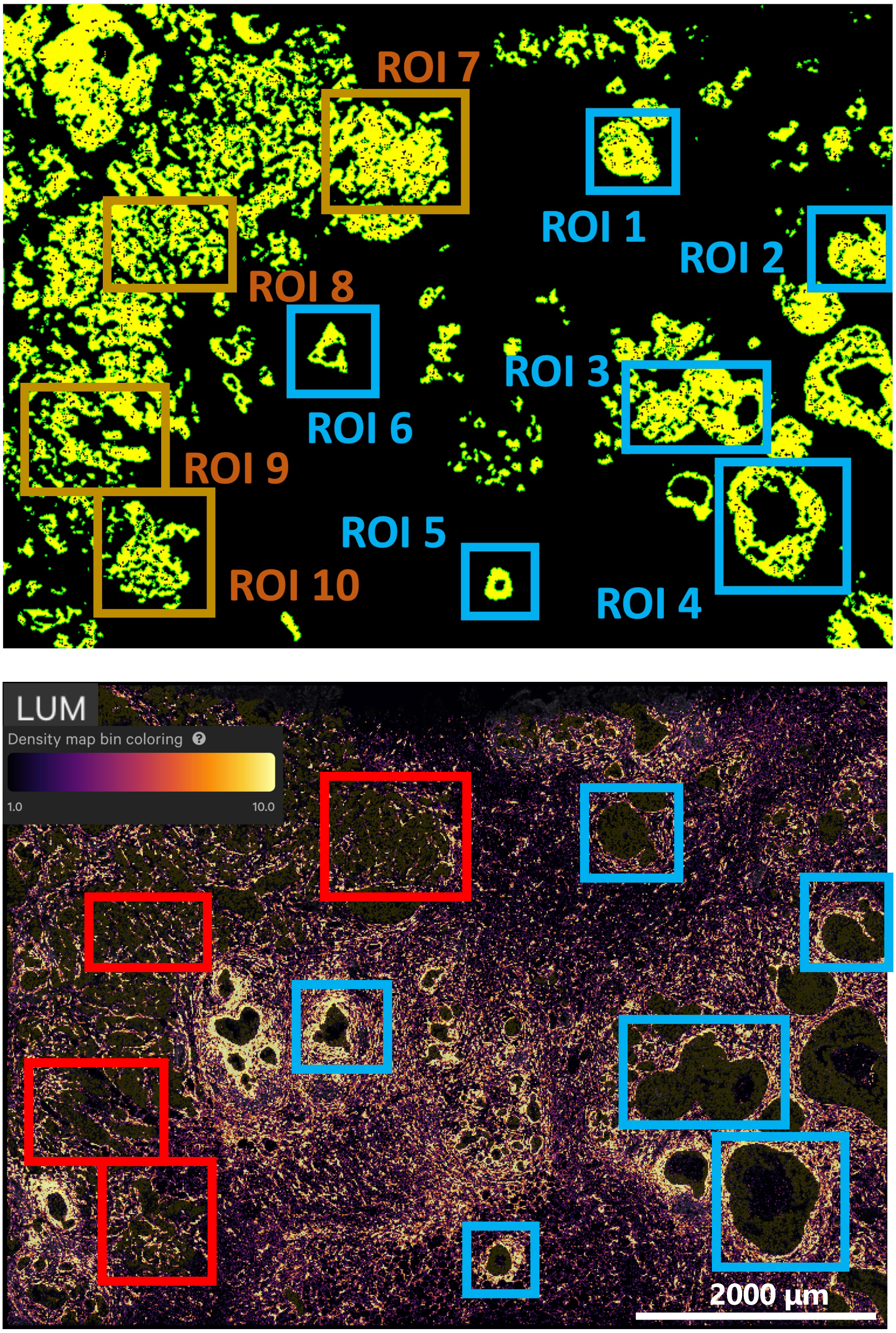


**Supplementary Fig. S27 Selected ROIs and spatial distribution of LUM.**
The upper panels show tumor regions identified by SpatialKnifeY (yellow) and the selected regions of interest (ROIs). The lower panels show the spatial expression of LUM, visualized using Xenium Explorer.

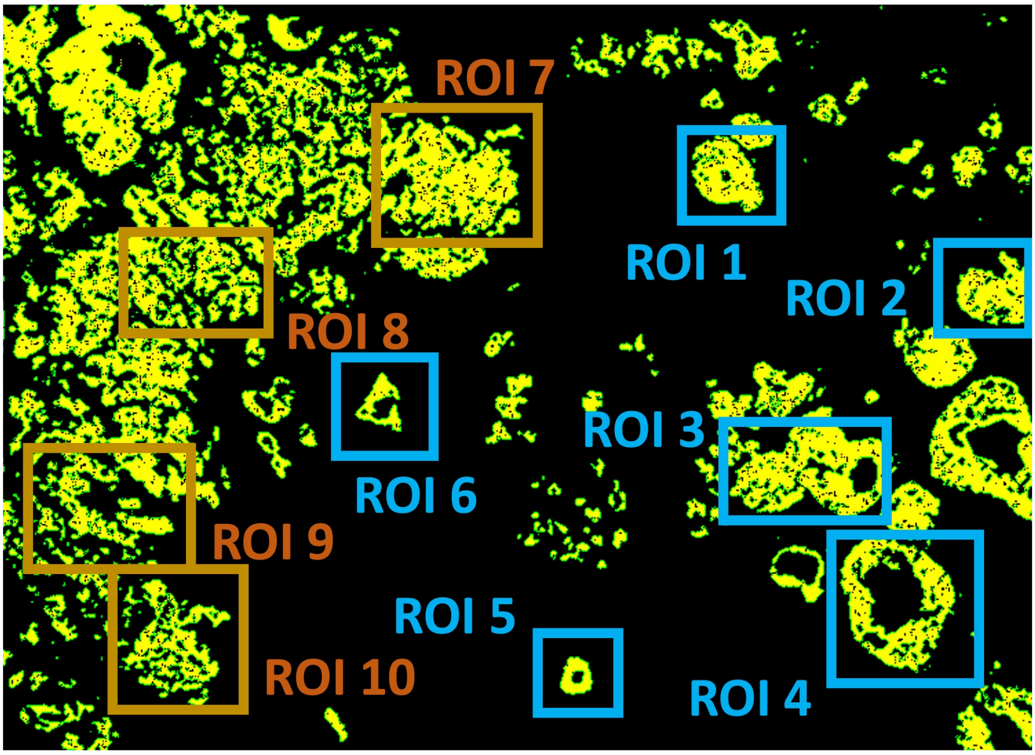


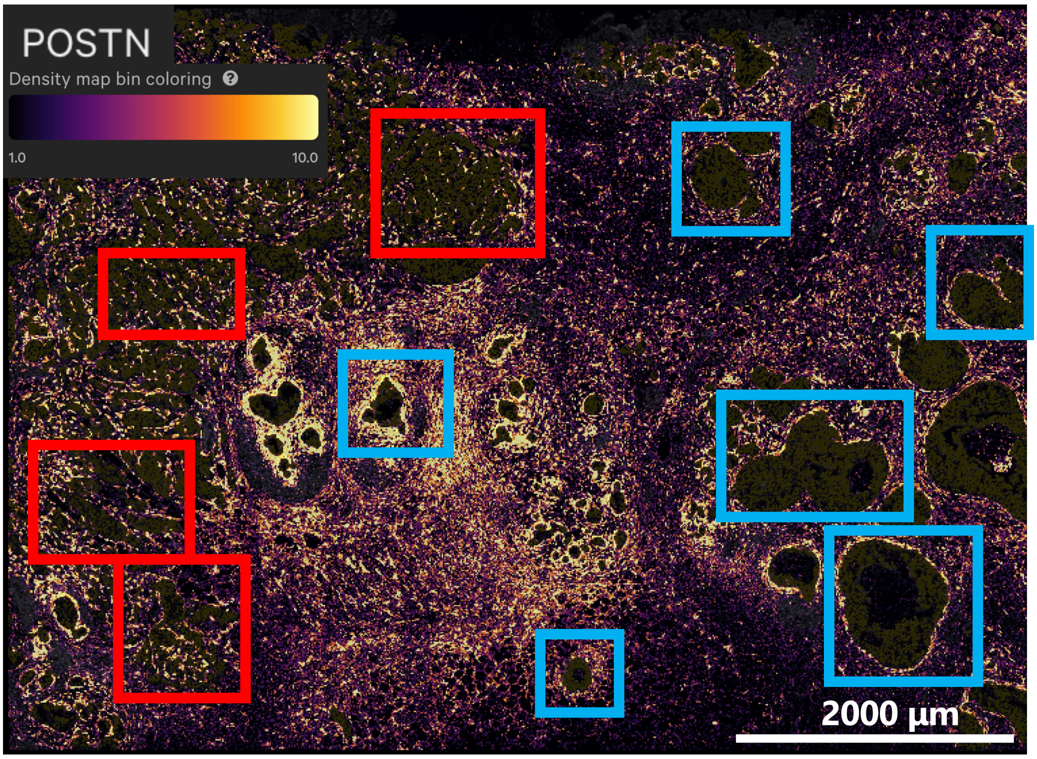


**Supplementary Fig. S28 Selected ROIs and spatial distribution of POSTN.**
The upper panels show tumor regions identified by SpatialKnifeY (yellow) and the selected regions of interest (ROIs). The lower panels show the spatial expression of POSTN, visualized using Xenium Explorer.

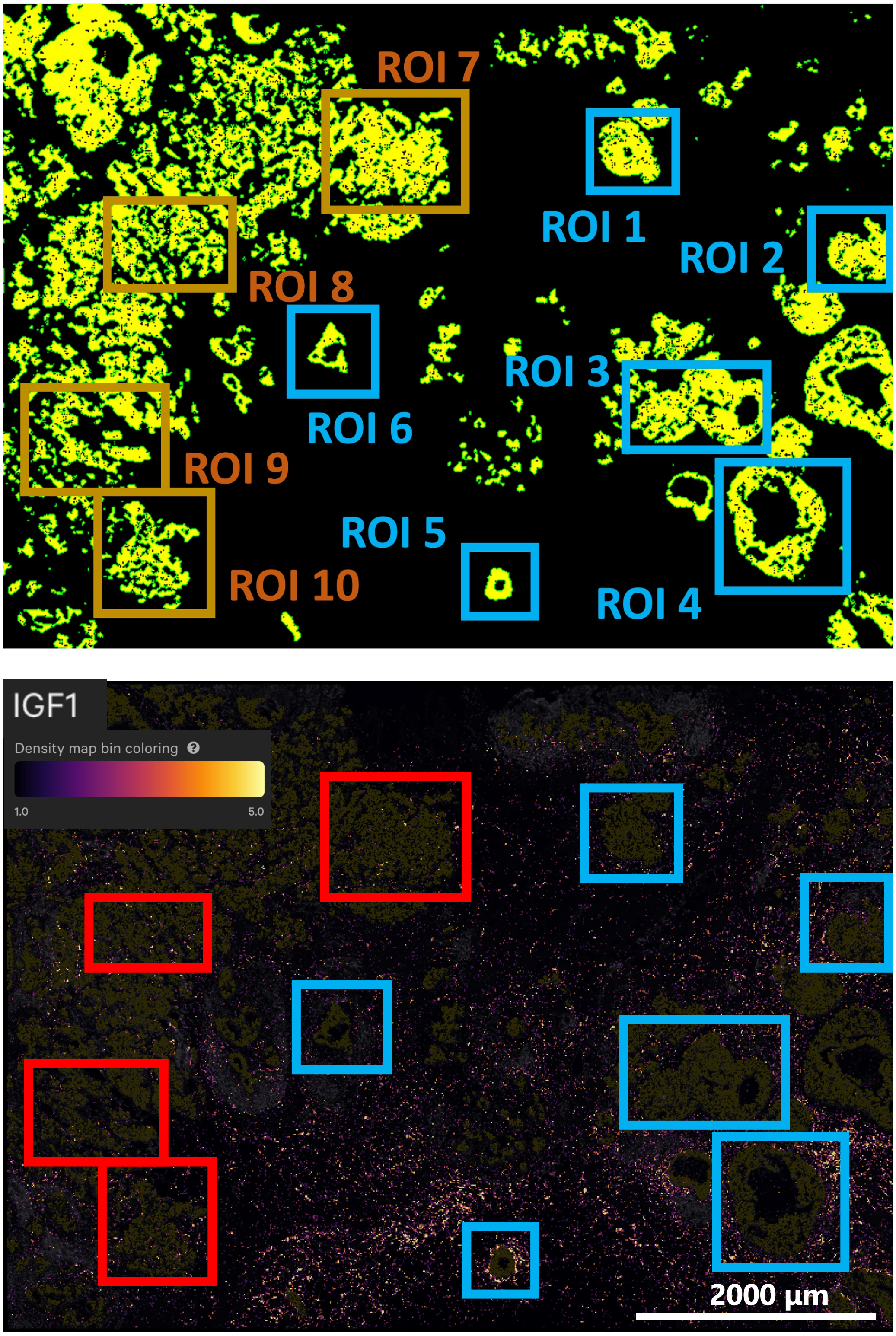


**Supplementary Fig. S29 Selected ROIs and spatial distribution of IGF1.**
The upper panels show tumor regions identified by SpatialKnifeY (yellow) and the selected regions of interest (ROIs). The lower panels show the spatial expression of IGF1, visualized using Xenium Explorer.

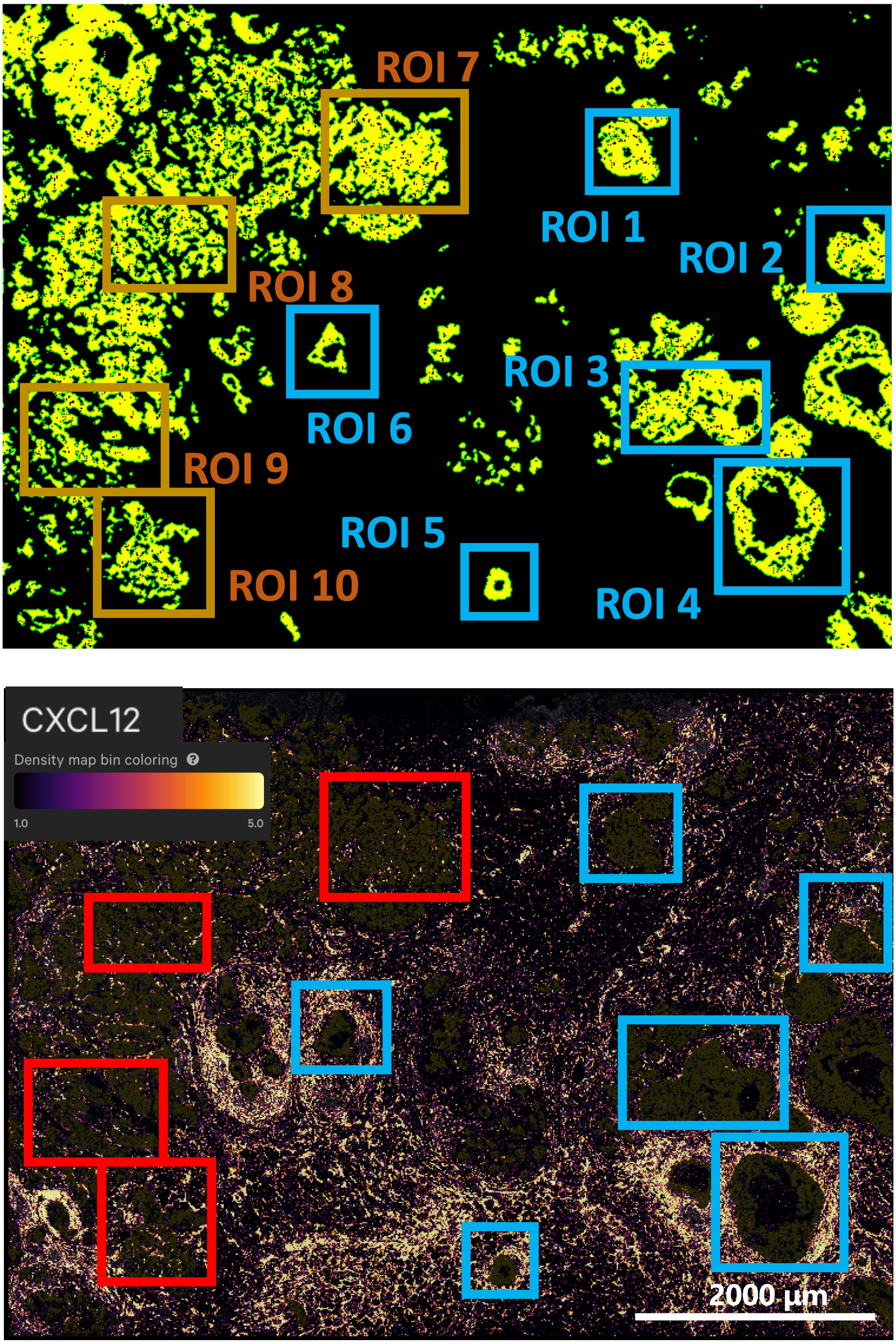


**Supplementary Fig. S30 Selected ROIs and spatial distribution of CXCL12.**
The upper panels show tumor regions identified by SpatialKnifeY (yellow) and the selected regions of interest (ROIs). The lower panels show the spatial expression of CXCL12, visualized using Xenium Explorer.

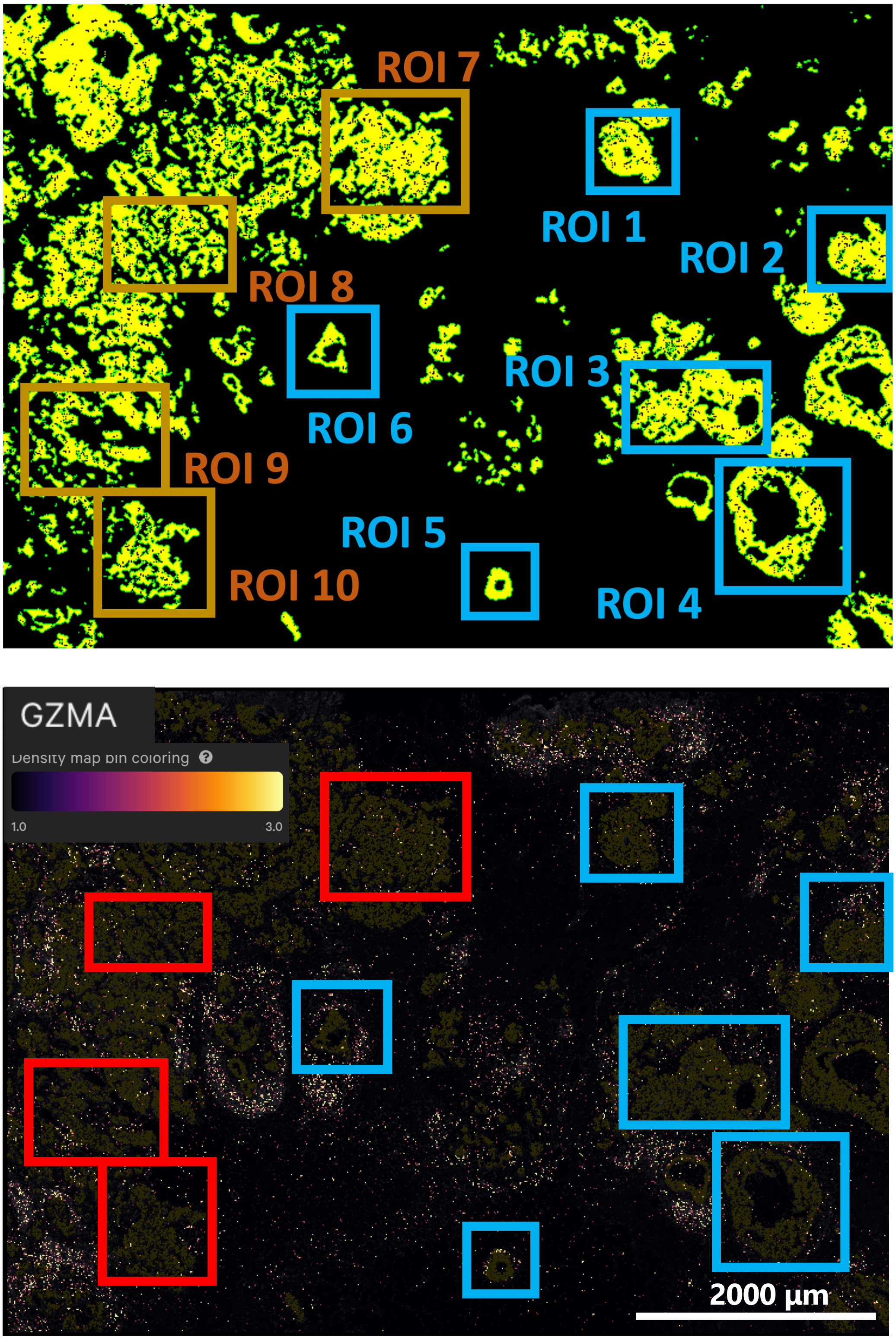


**Supplementary Fig. S31 Selected ROIs and spatial distribution of GZMA.**
The upper panels show tumor regions identified by SpatialKnifeY (yellow) and the selected regions of interest (ROIs). The lower panels show the spatial expression of GZMA, visualized using Xenium Explorer.

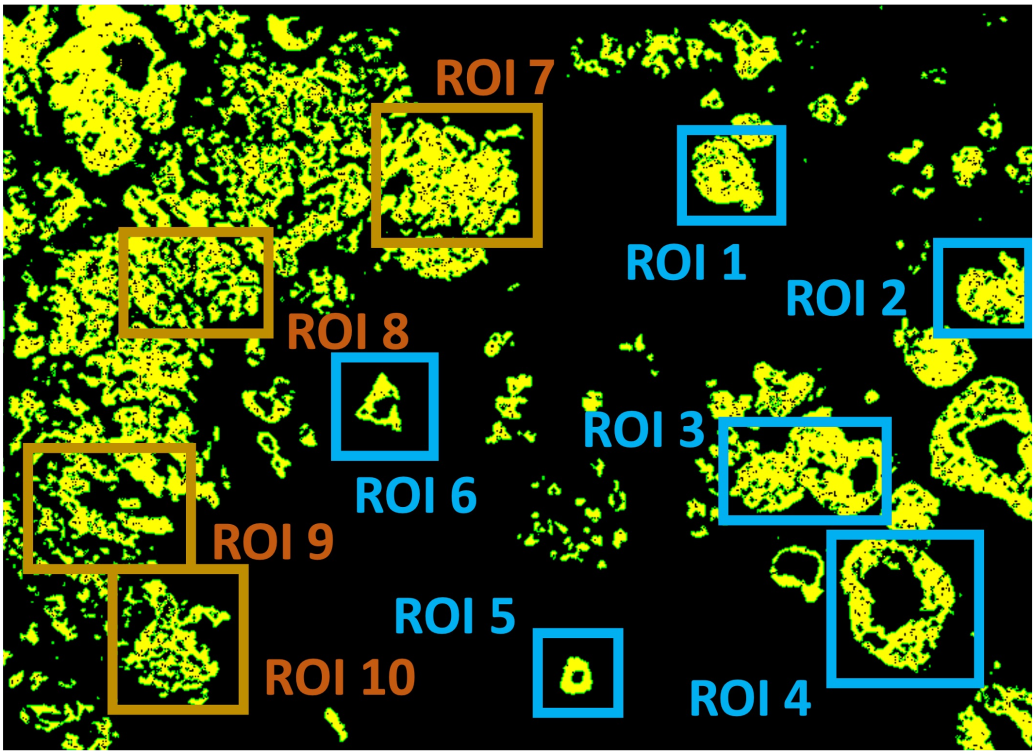


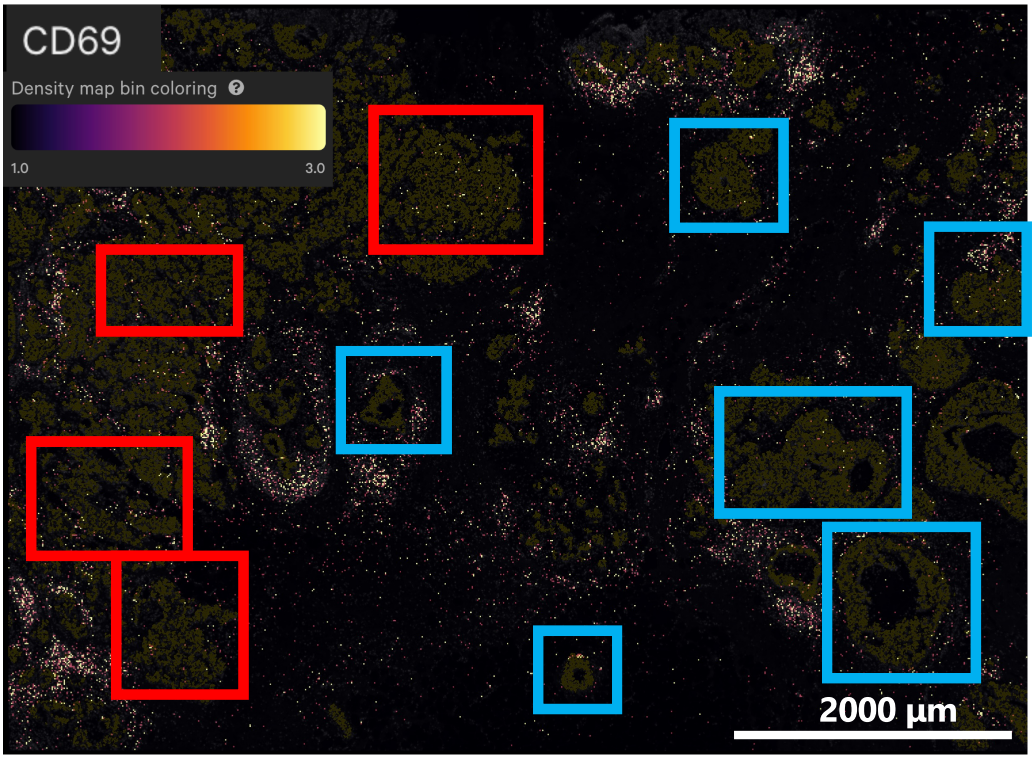


**Supplementary Fig. S32 Selected ROIs and spatial distribution of CD69.**
The upper panels show tumor regions identified by SpatialKnifeY (yellow) and the selected regions of interest (ROIs). The lower panels show the spatial expression of CD69, visualized using Xenium Explorer.

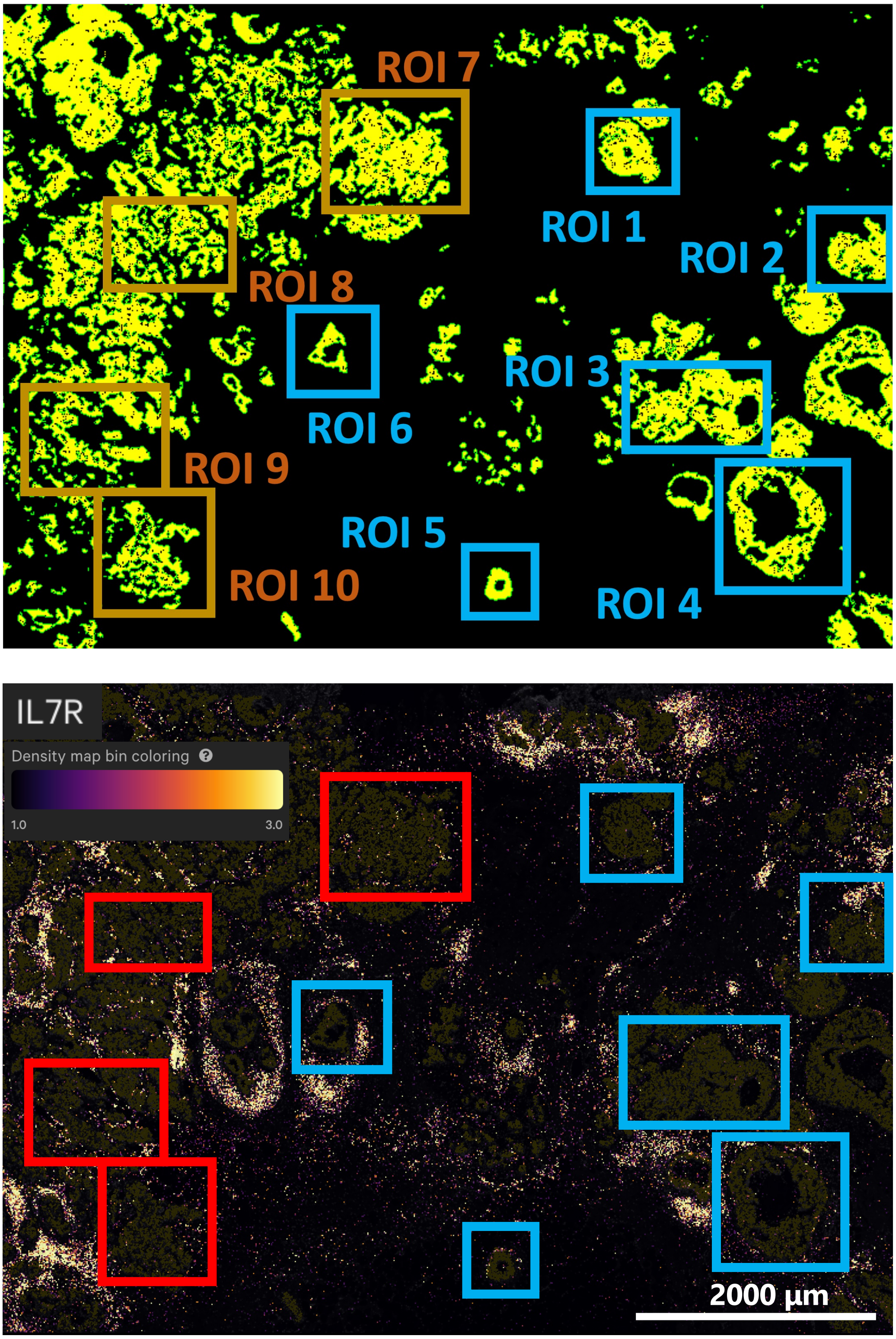


**Supplementary Fig. S33 Selected ROIs and spatial distribution of IL7R.**
The upper panels show tumor regions identified by SpatialKnifeY (yellow) and the selected regions of interest (ROIs). The lower panels show the spatial expression of IL7R, visualized using Xenium Explorer.

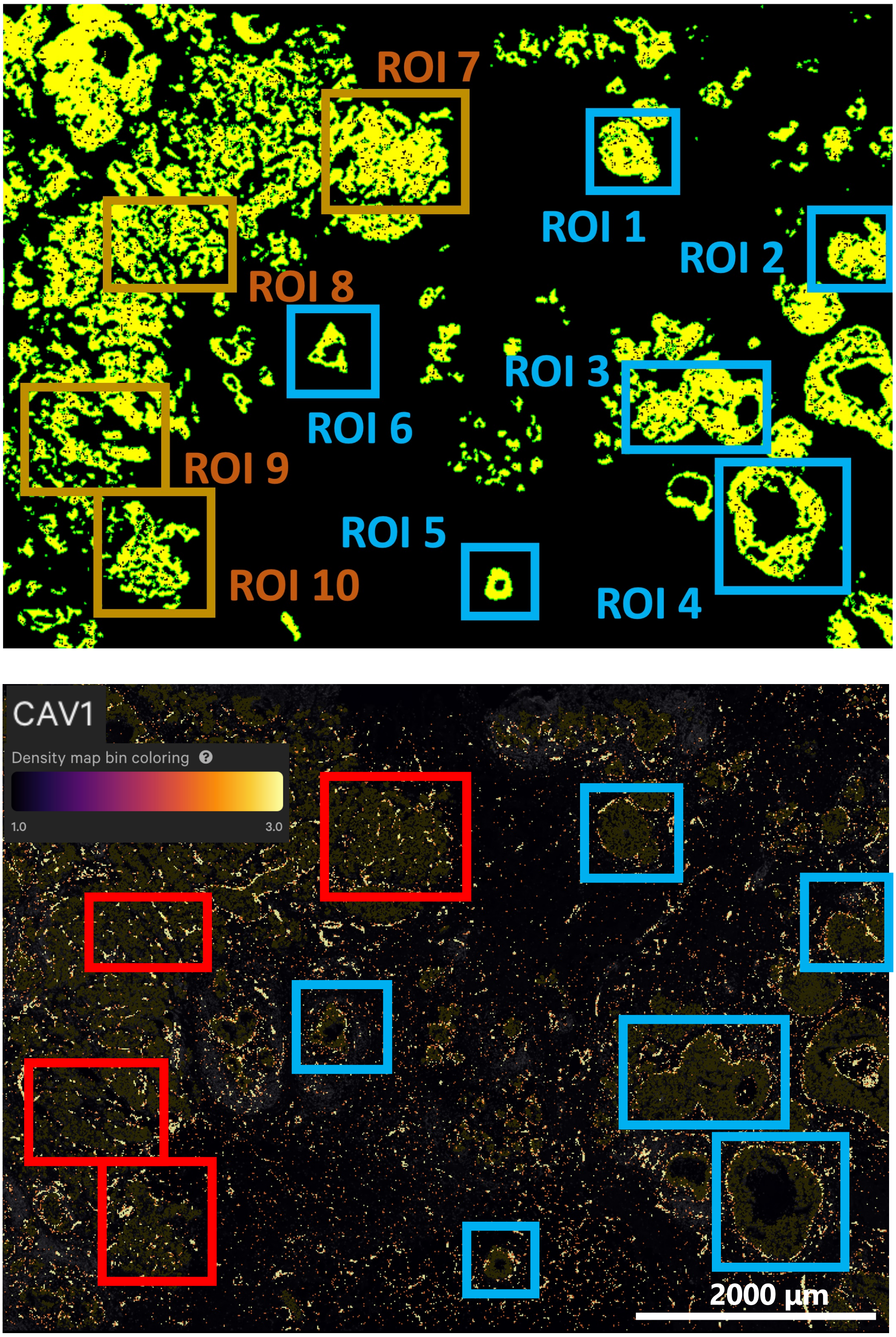


**Supplementary Fig. S34 Selected ROIs and spatial distribution of CAV1.**
The upper panels show tumor regions identified by SpatialKnifeY (yellow) and the selected regions of interest (ROIs). The lower panels show the spatial expression of CAV1, visualized using Xenium Explorer.

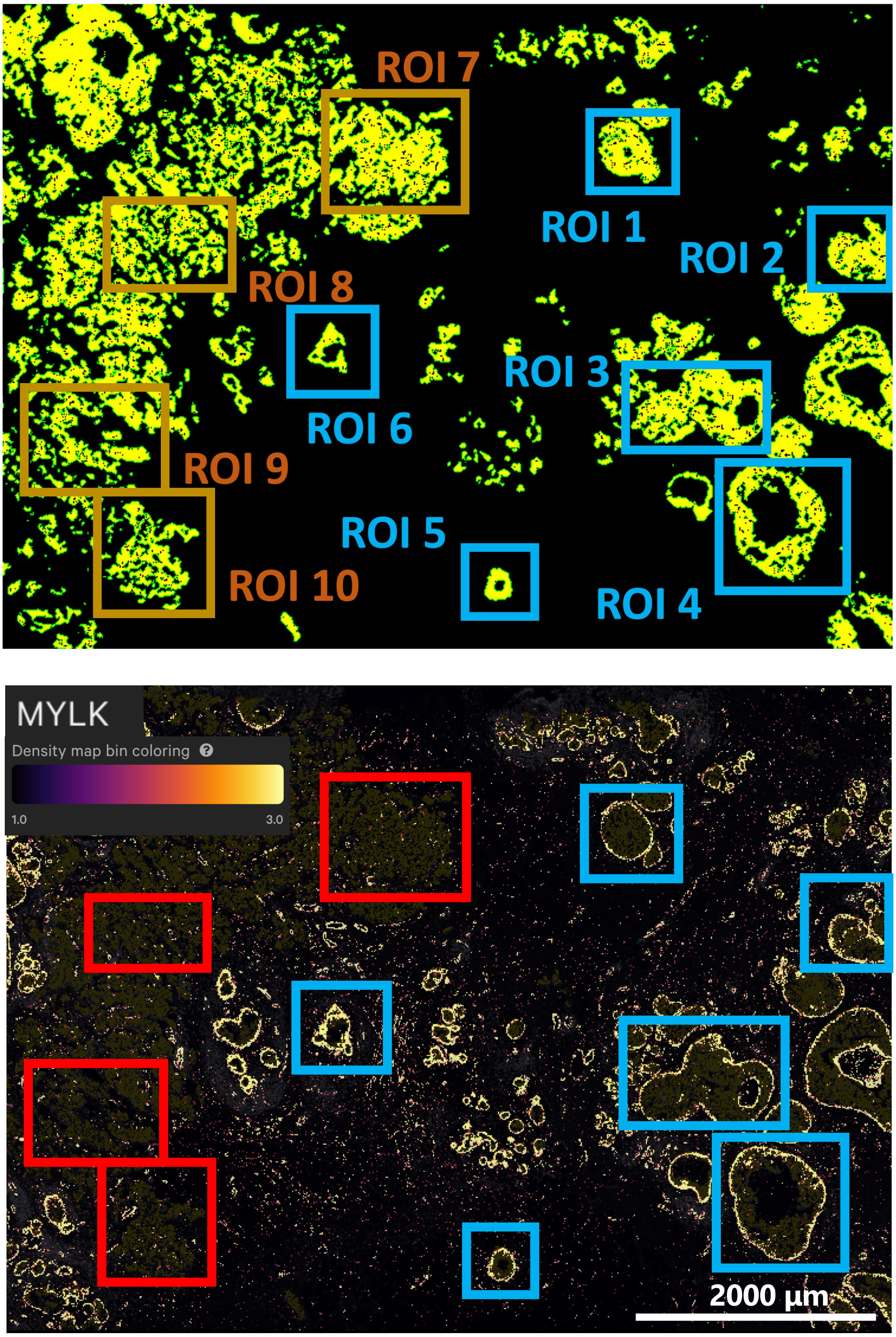


**Supplementary Fig. S35 Selected ROIs and spatial distribution of MYLK.**
The upper panels show tumor regions identified by SpatialKnifeY (yellow) and the selected regions of interest (ROIs). The lower panels show the spatial expression of MYLK, visualized using Xenium Explorer.

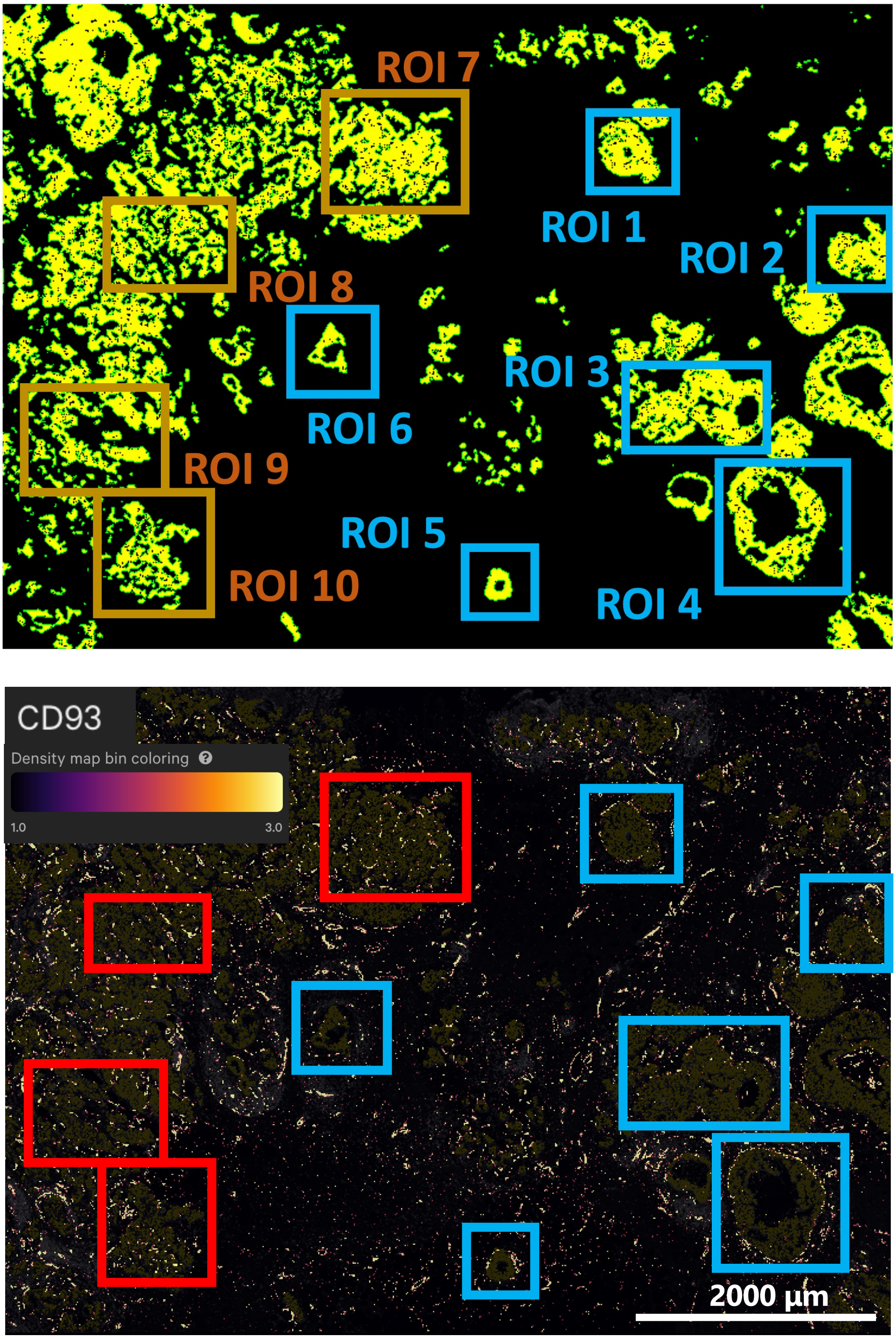


**Supplementary Fig. S36 Selected ROIs and spatial distribution of CD93.**
The upper panels show tumor regions identified by SpatialKnifeY (yellow) and the selected regions of interest (ROIs). The lower panels show the spatial expression of CD93, visualized using Xenium Explorer.

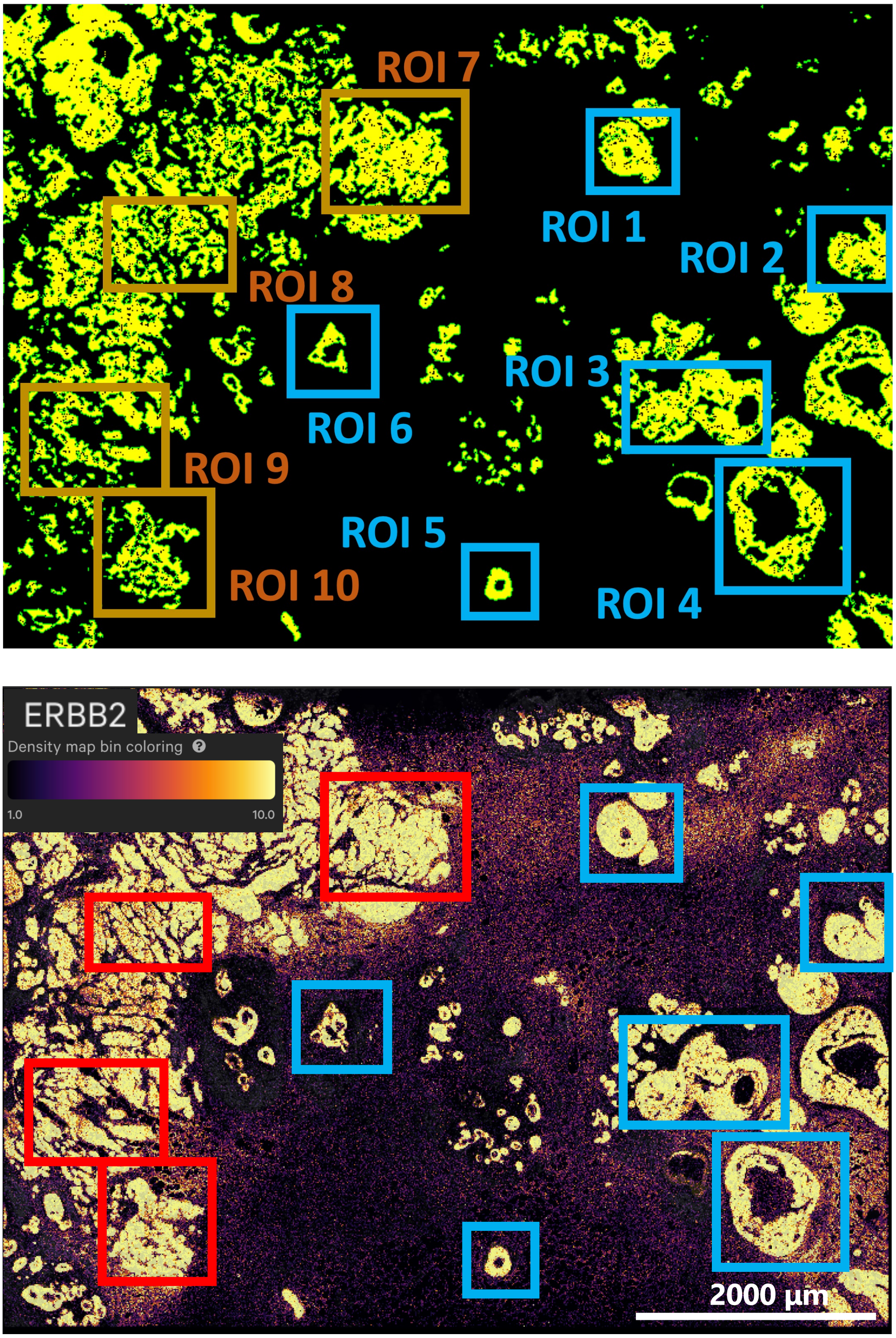


**Supplementary Fig. S37 Selected ROIs and spatial distribution of ERBB2.**
The upper panels show tumor regions identified by SpatialKnifeY (yellow) and the selected regions of interest (ROIs). The lower panels show the spatial expression of ERBB2, visualized using Xenium Explorer.

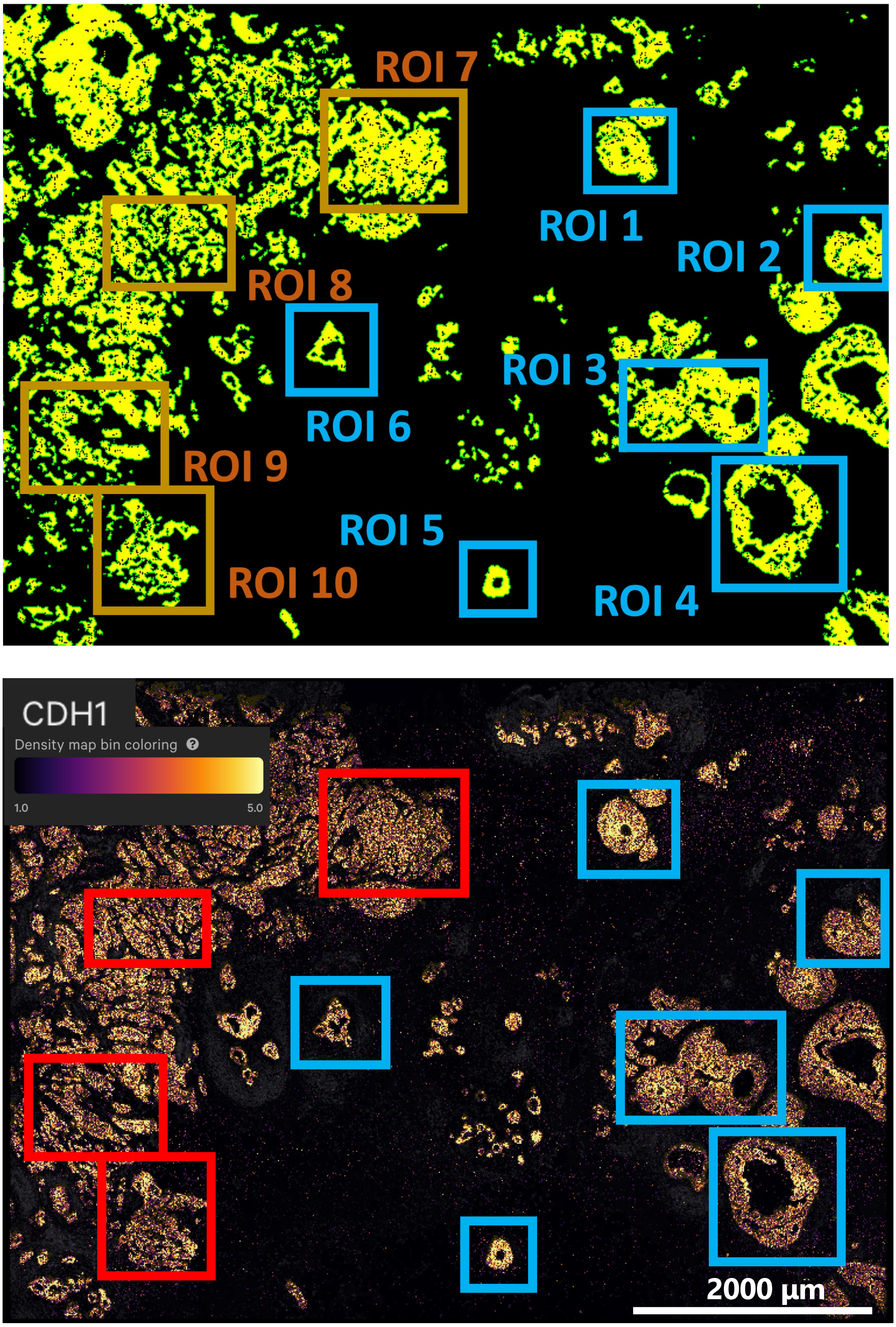


**Supplementary Fig. S38 Selected ROIs and spatial distribution of CDH1.**
The upper panels show tumor regions identified by SpatialKnifeY (yellow) and the selected regions of interest (ROIs). The lower panels show the spatial expression of CDH1, visualized using Xenium Explorer.

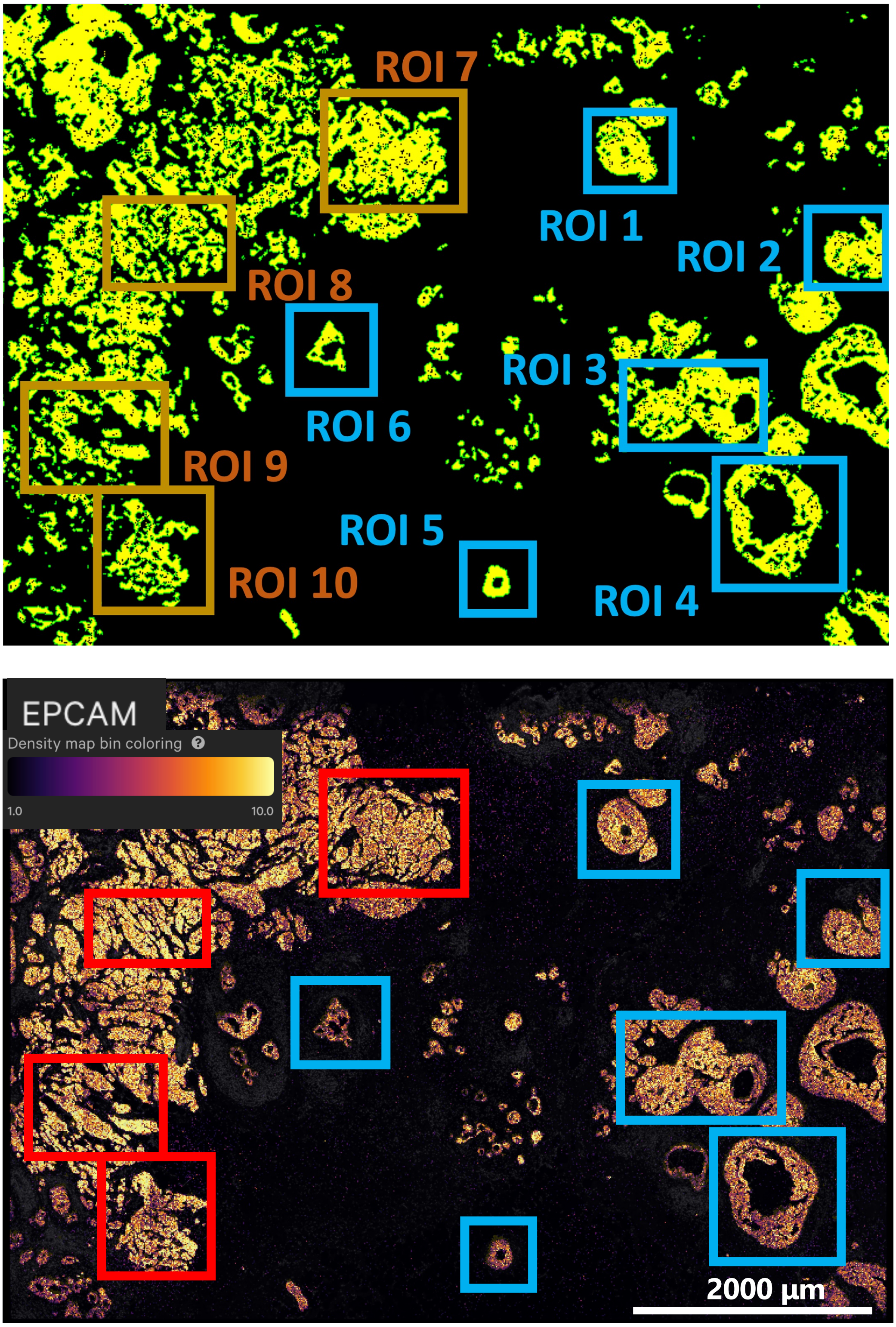


**Supplementary Fig. S39 Selected ROIs and spatial distribution of EPCAM.**
The upper panels show tumor regions identified by SpatialKnifeY (yellow) and the selected regions of interest (ROIs). The lower panels show the spatial expression of EPCAM, visualized using Xenium Explorer.
